# Regulation of the 20S proteasome by a novel family of inhibitory proteins

**DOI:** 10.1101/617415

**Authors:** Maya A Olshina, Fanindra Kumar Deshmukh, Galina Arkind, Irit Fainer, Mark Taranavsky, Daniel Hayat, Shifra Ben-Dor, Gili Ben-Nissan, Michal Sharon

**Affiliations:** Department of Biomolecular Sciences, Weizmann Institute of Science, Rehovot 7610001, Israel; Bioinformatics and Biological Computing Unit, Weizmann Institute of Science, Rehovot 7610001, Israel

## Abstract

The protein degradation machinery plays a critical role in the maintenance of cellular homeostasis, preventing the accumulation of damaged or misfolded proteins and controlling the levels of regulatory proteins. The 20S proteasome degradation machinery is able to cleave any protein with a partially unfolded region, however uncontrolled degradation of the myriad of potential substrates is improbable. Thus, there must exist a regulatory mechanism to control 20S proteasome mediated degradation. Here we have discovered a family of 20S proteasome regulators, named Catalytic Core Regulators (CCRs). They coordinate the function of the 20S proteasome and are involved in the oxidative stress response via Nrf2. The CCRs organize into a feed-forward loop regulatory circuit, with some members stabilizing Nrf2, others being induced by Nrf2, and all of them inhibiting the 20S proteasome. This provides a fine-tuned mechanism to carefully modulate the 20S proteasome, ensuring its proper functioning by controlling the degradative flux.

## Introduction

The ability to maintain cellular homeostasis, while being able to rapidly respond to environmental stimuli and stressors, requires a cell to coordinate multiple complex pathways to ensure ongoing cell health and survival (Labbadia & Morimoto, 2015). In order to meet the fluctuating needs of the cell, turnover of proteins is necessary to adapt the composition of the proteome as required. The major process through which proteins are degraded involves the ubiquitin-proteasome system, a complicated network of enzymes and degradation machinery that breaks down proteins marked for destruction (Ciechanover & Schwartz, 2009, Finley, 2009, Goldberg, 2003). This pathway begins with the conjugation of ubiquitin chains to the protein to be degraded via an enzymatic cascade, followed by recognition of the ubiquitin by the 26S proteasome complex (Saeki, 2017). The 26S proteasome is comprised of a 19S regulatory particle, which recognizes ubiquitin tagged substrates, and a 20S catalytic core particle, in which substrates are degraded via breakage of peptide bonds. The ATPase activity of several subunits of the 19S provides the energy required to unfold the protein substrate and translocate it into the central pore of the 20S, where proteolysis occurs by the activity of its catalytic subunits.

While protein degradation via the ubiquitin-26S proteasome pathway is the predominant mechanism used by the cell to degrade proteins, it has been well established that the 20S catalytic core of the proteasome is capable of operating independently of the 19S regulatory particle (Baugh, Viktorova et al., 2009, Ben-Nissan & Sharon, 2014, Davies, 2001, Olshina, Ben-Nissan et al., 2017, Pickering & Davies, 2012). The 20S proteasome can degrade protein substrates in an ubiquitin and ATP-independent manner, by recognizing unfolded or unstructured regions within its substrates, as opposed to the specificity of a ubiquitin tag as required by the 26S proteasome (Erales & Coffino, 2014, Sanchez-Lanzas & Castano, 2014). Proteins can acquire unstructured regions as a result of mutation, aging or damage, which can occur under stress conditions such as oxidative stress (Pickering & Davies, 2012). Alternatively, proteins can contain intrinsically disordered regions (IDRs), for example the cell cycle regulators p21 and p27, and tumor suppressors p53 and p73 (Dyson & Wright, 2005, Iakoucheva, Brown et al., 2002, Van Der Lee, Buljan et al., 2014, Yoon, Mitrea et al., 2012). Beyond complete degradation of unstructured protein substrates, the 20S proteasome has also been shown to be responsible for the post-translational processing of certain proteins, generating proteolytic products that display unique roles and functionality compared with their parent proteins (Baugh & Pilipenko, 2004, Moorthy, Savinova et al., 2006, Morozov, Astakhova et al., 2017, Olshina et al., 2017, Solomon, Bräuning et al., 2017).

Notably, it has been demonstrated that 20% of all cellular proteins are degraded in an ubiquitin-independent manner by the 20S proteasome (Baugh et al., 2009), and analysis of the human genome has indicated that almost half of all proteins are predicted to contain disordered segments (Dyson & Wright, 2005, Van Der Lee et al., 2014). Thus, the theoretical substrate pool for the 20S proteasome is substantial, even in unstressed cells without widespread damage to proteins. In addition, recent studies have quantified the proportions of the various proteasome complexes across multiple cell types, revealing that the free 20S proteasome is consistently the most abundant form of the proteasome in cells (Fabre, Lambour et al., 2013, Fabre, Lambour et al., 2014). Furthermore, under conditions of oxidative stress, there is *de novo* synthesis of 20S proteasome subunits and disassembly of the 26S proteasome into its 20S and 19S components, increasing the total amount of free 20S proteasome in the cell (Aiken, Wang et al., 2011, Grune, Catalgol et al., 2011, Wang, Chemmama et al., 2017). The 20S proteasome remains catalytically active and has been implicated in the degradation of oxidatively damaged proteins, preventing them from accumulating and causing cytotoxicity and cell death (Pickering & Davies, 2012). Oxidative stress also leads to the inactivation of ubiquitin conjugating enzymes, disrupting the ubiquitination cascade and thus reducing the flux of proteins through the 26S proteasome pathway (Shang & Taylor, 2011).

The relatively large proportion of free 20S proteasomes, in particular under conditions of oxidative stress, is therefore theoretically capable of degrading a significant amount of proteins within the cell, both under normal and stress conditions. Given the broad substrate landscape of the 20S proteasome, which encompasses not only damaged proteins but also important regulatory proteins, it is unlikely that 20S proteasome mediated degradation persists in an unregulated manner, as critical imbalances may occur. In addition, the significant increase in damaged proteins during stress that become potentially pathological and prone to aggregation and toxicity and require degradation, could overload the 20S proteasomes leading to proteasome clogging and consequently dysfunction. Therefore, it is reasonable to assume that the flux of substrate proteins through the 20S proteasome pathway will be tightly regulated (Figure 1).

**Figure 1:**
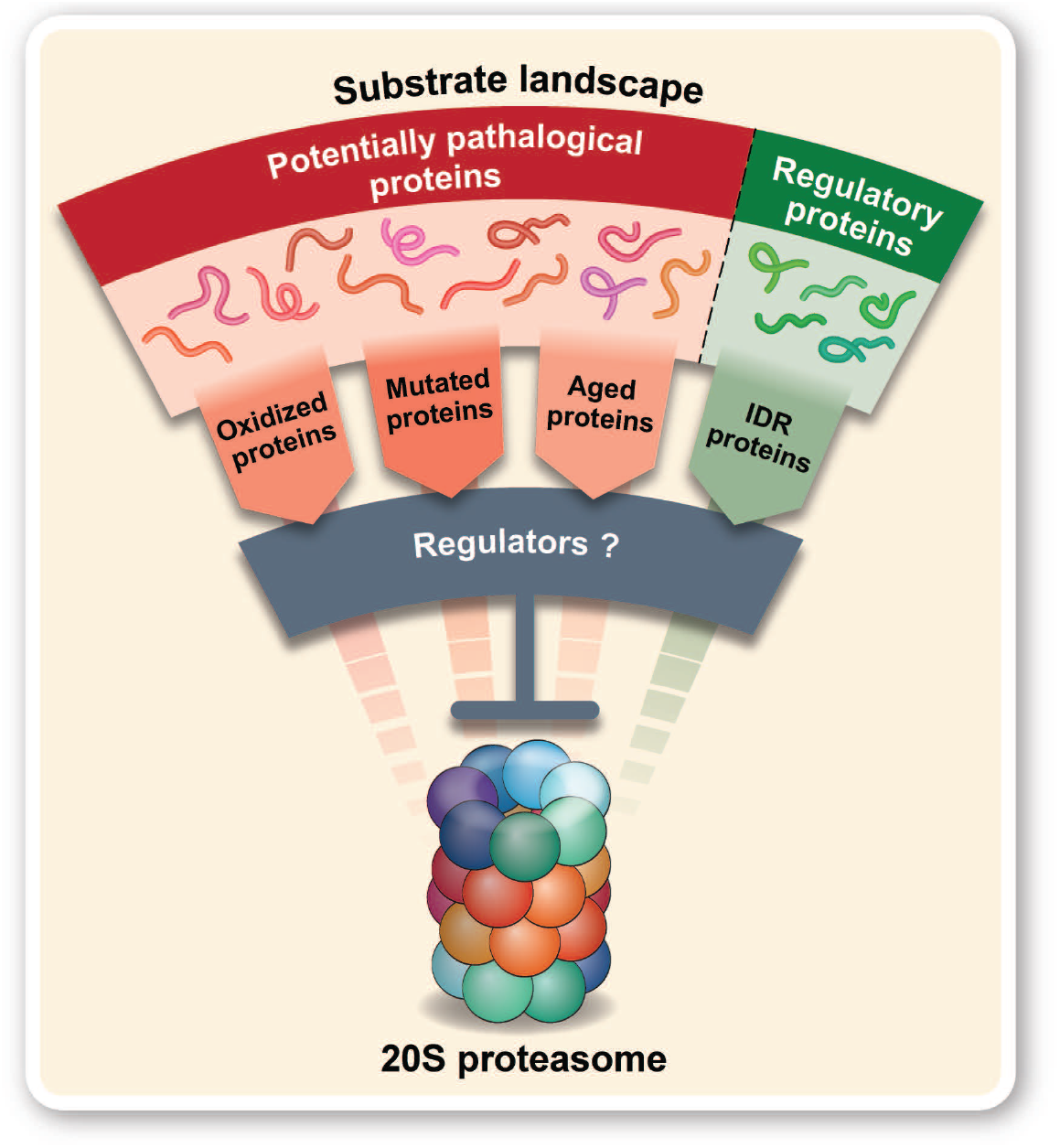
The flux of substrates into the 20S proteasome must be a regulated process. Two main groups of substrates are susceptible to 20S degradation. The first consists of proteins that have lost their native structure due to aging, mutations or oxidative damage. These proteins are prone to aggregation and may lead to cytotoxicity, and, therefore, should be rapidly removed to prevent cell malfunctions that lead to pathologies, such as neurodegenerative disorders. The second group comprises substrates with unfolded regions as an intrinsic feature of the proteins themselves. Many key regulatory proteins that belong to this group have been shown to be substrates of the 20S proteasome. The regulation of the flux of substrates that enter into the 20S could prevent proteasome clogging and excessive degradation of essential regulatory proteins.

While the presence of a regulatory mechanism to control this degradation process is highly likely, to date only two 20S proteasome regulators have been identified; NQO1 (Asher, Tsvetkov et al., 2005, Moscovitz, Tsvetkov et al., 2012) and DJ-1 (Moscovitz, Ben-Nissan et al., 2015). These proteins share multiple similarities in structure and function, such as the presence of a Rossman fold, their role in the cellular response to oxidative stress and the ability to ‘moonlight’ as 20S proteasome inhibitors while being enzymatically active in other cellular pathways (Moscovitz et al., 2015). Given the cellular implications of improper regulation of the 20S proteasome, and the fact the DJ-1 was explored as a potential 20S proteasome regulator based on its similarities to NQO1, we hypothesized that there may exist a broader family of proteins that exhibit these features, and thus may also be able to regulate the 20S proteasome.

Here we performed a bioinformatics screen to identify other proteins with sequence and structural similarities to DJ-1 and NQO1. This led to the discovery of a family of 17 relatively small proteins of 20-30 kDa, which we named Catalytic Core Regulators (CCRs) that oversee 20S proteasome activity. Of the ten shortlisted proteins that were identified and characterized, all were able to inhibit 20S proteasome degradation of known substrates, both *in vitro* and in cells. These protein regulators were able to specifically bind to the 20S proteasome, but not the 26S proteasome, and affect protein degradation. In addition, we demonstrate that the CCRs organize into a feed forward regulation circuit involving the master regulator of the oxidative stress response, Nrf2. Certain CCRs influence the stability of Nrf2, which subsequently upregulates the expression of the other CCRs, leading to an overall dampening of 20S proteasome mediated degradation of unfolded protein substrates within the cell. Overall, our results suggest that 20S proteasome mediated degradation is not a simple and random process, but rather a highly CCR- regulated and coordinated mechanism.

## Results

### DJ-1 inhibition of the 20S proteasome is conserved across evolution

Previously, we discovered that DJ-1, the Parkinson’s disease related protein, is a regulator of the 20S proteasome (Moscovitz et al., 2015). To determine if the inhibitory capacity of DJ-1 is evolutionarily conserved, we began by analyzing its ability to inhibit 20S proteasomes purified from three different sources; mammals, yeast and archaea (Figure 2A). The IDR containing model substrate α-synuclein (α-syn) was used to monitor 20S proteasome mediated degradation over time. Inhibition of the human and yeast 20S proteasomes occurred to a similar degree, and despite the vast evolutionary distance, human DJ-1 was also able to reduce the rate of degradation of the archaeal 20S proteasome. The reciprocal experiment was also performed, in which DJ-1 homologues from mammals, yeast and archaea were tested for their ability to inhibit the mammalian 20S proteasome (Figure 2B). All three DJ-1 homologues were able to inhibit the 20S proteasome, reducing the rate of degradation of α-syn over the course of the experiment. Taken together, these results indicate that the inhibition of the 20S proteasome by DJ-1 is conserved across evolution, highlighting the essentiality of this cellular process.

**Figure 2:**
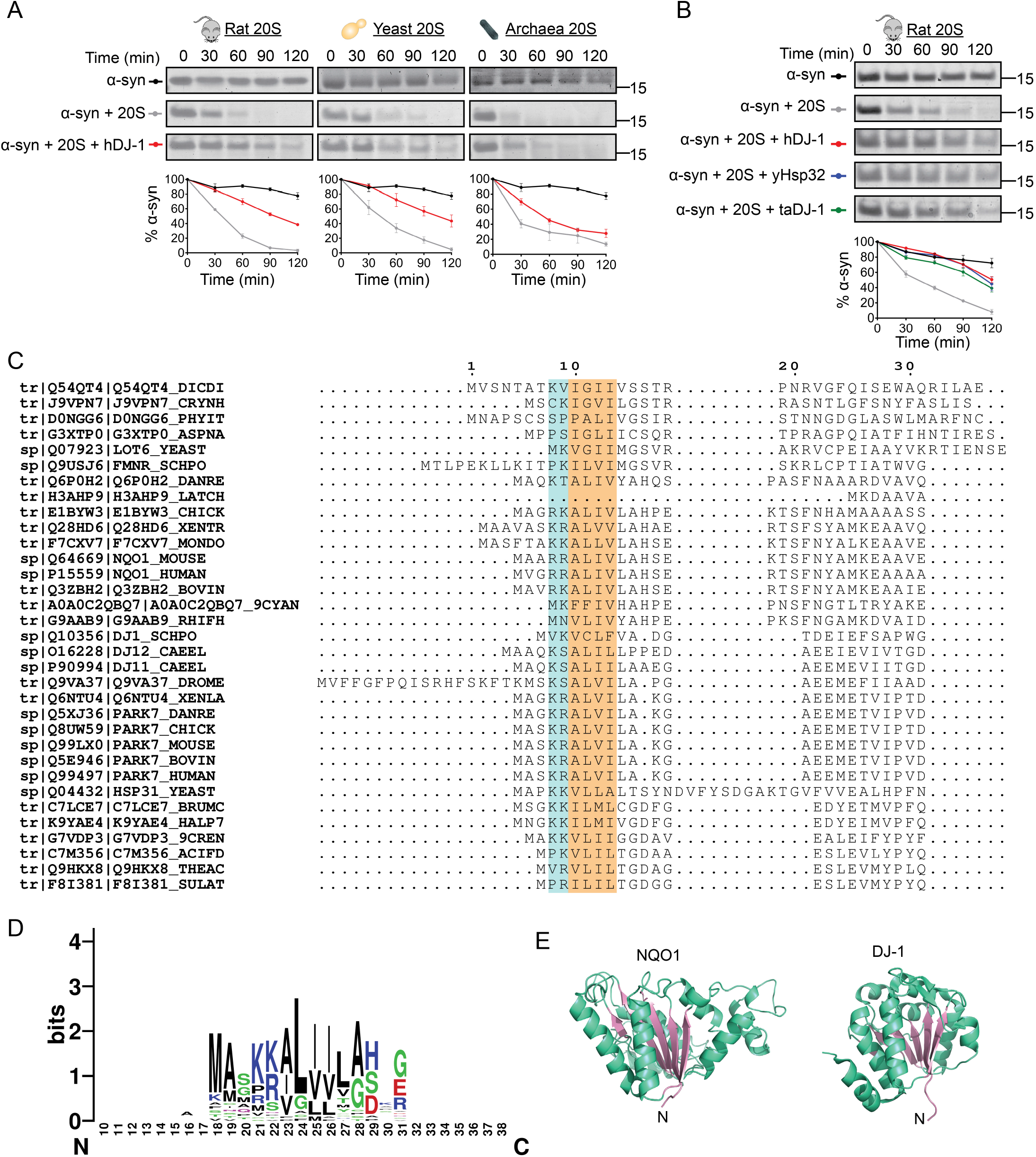
Evolutionary conservation of the regulatory activity of DJ-1 on the 20S proteasome and the discovery of a conserved N-terminal motif. (A) *In vitro* degradation assays using model substrate α-syn were performed using mammalian DJ-1 (*H. sapiens*, hDJ-1) with 20S proteasomes from three different sources; mammals (*R. norvegicus*, r20S), yeast (*S. cerevisiae*, y20S) or archaea (*T. acidophilum*, ta20S). (B) *In vitro* degradation assays were performed using 20S proteasomes from mammals (r20S) with DJ-1 homologues from three different sources (mammals (hDJ-1), yeast (yHsp32) or archaea (taDJ-1). Representative coomassie stained SDS-PAGE gels with quantification of three independent experiments are shown. The three graphs in (A) were derived from the same set of experiments, therefore the quantification of α-syn is identical in the graphs. The results are displayed separately for ease of viewing the inhibitory effect of hDJ-1 on each 20S proteasome. Error bars represent S.E.M. (C) Multiple sequence alignment (MSA) of NQO1 and DJ-1 homologues reveals conservation of two positive residues (highlighted in blue) followed by at least four hydrophobic residues (highlighted in orange) near the N-terminus of the proteins. (D) Weblogo representation of the amino acid conservation near the N-terminus of the MSA. (E) Structures of NQO1 (1D4A.pdb) and DJ-1 (1UCF.pdb) with the central β-sheet of the Rossman fold highlighted in pink, and the α-helices in green.

### Bioinformatic screen identified a new family of Catalytic Core Regulators

The ability of DJ-1 to inhibit 20S proteasomes from such evolutionarily distant species not only indicates that this process evolved early, but it also provides the basis for searching for additional proteins with similar properties. In addition, a homologue of NQO1 from yeast, Lot6, was also shown to interact with and inhibit the 20S proteasome, further supporting the view that this is a conserved process (Sollner, Schober et al., 2009). Therefore, a bioinformatics approach was used to reveal the sequence and structural similarities of NQO1 and DJ-1, with the rationale that identified features would then be exploited to search for new 20S proteasome regulatory candidates. As a first step, a multiple sequence alignment of NQO1 and DJ-1 homologues from a wide variety of species was performed (Figure 2C), revealing a conserved region towards the N-terminus of the proteins. This motif consists of two positively charged residues (K or R) followed by at least four hydrophobic residues (A, V, I or L) (Figure 2D). Both NQO1 and DJ-1 adopt a Rossman fold, composed of an extended parallel β-sheet with α-helices surrounding both faces to produce a three-layered α/β/α sandwich (Figure 2E). Therefore, the search for new 20S proteasome regulatory candidates was restricted to proteins which have been classified as Rossman fold containing proteins, comprise the conserved N-terminal motif, and are less than 100kDa in size.

The search yielded 17 proteins, including NQO1 and DJ-1, herein referred to as Catalytic Core Regulators (CCRs) (Table 1). Interestingly, all of these proteins range in size from 20-30kDa, and examination of the literature indicated that many of them are known to be connected to the oxidative stress response in cells, with links to Nrf2 (references listed in Table 1). Included among the potential CCRs are several well characterized enzymes, such as the quinone reductase enzyme NRH:quinone oxidoreductase 2 (NQO2), which is in the same enzyme family as NQO1, carbonyl reductase 3 (CBR3) and 15-hydroxyprostaglandin dehydrogenase (PGDH). Multiple members of the Ras superfamily of proteins are represented, such as NRas, KRas and HRas, as well as Rho and Rap family proteins. Interestingly, analysis of the tissue wide protein expression levels of 11 of these putative CCRs demonstrate that some of them display specificity to certain tissues, such as RBBP9 in the oral epithelium and PGDH in the lung, while DJ-1 and RhoA display a significantly more widespread expression profile than that of the other proteins (Supplementary Figure 1). This hints towards variation in the activities of these proteins, possibly indicating additional significance to the roles of DJ-1 and RhoA.

**Table 1:**
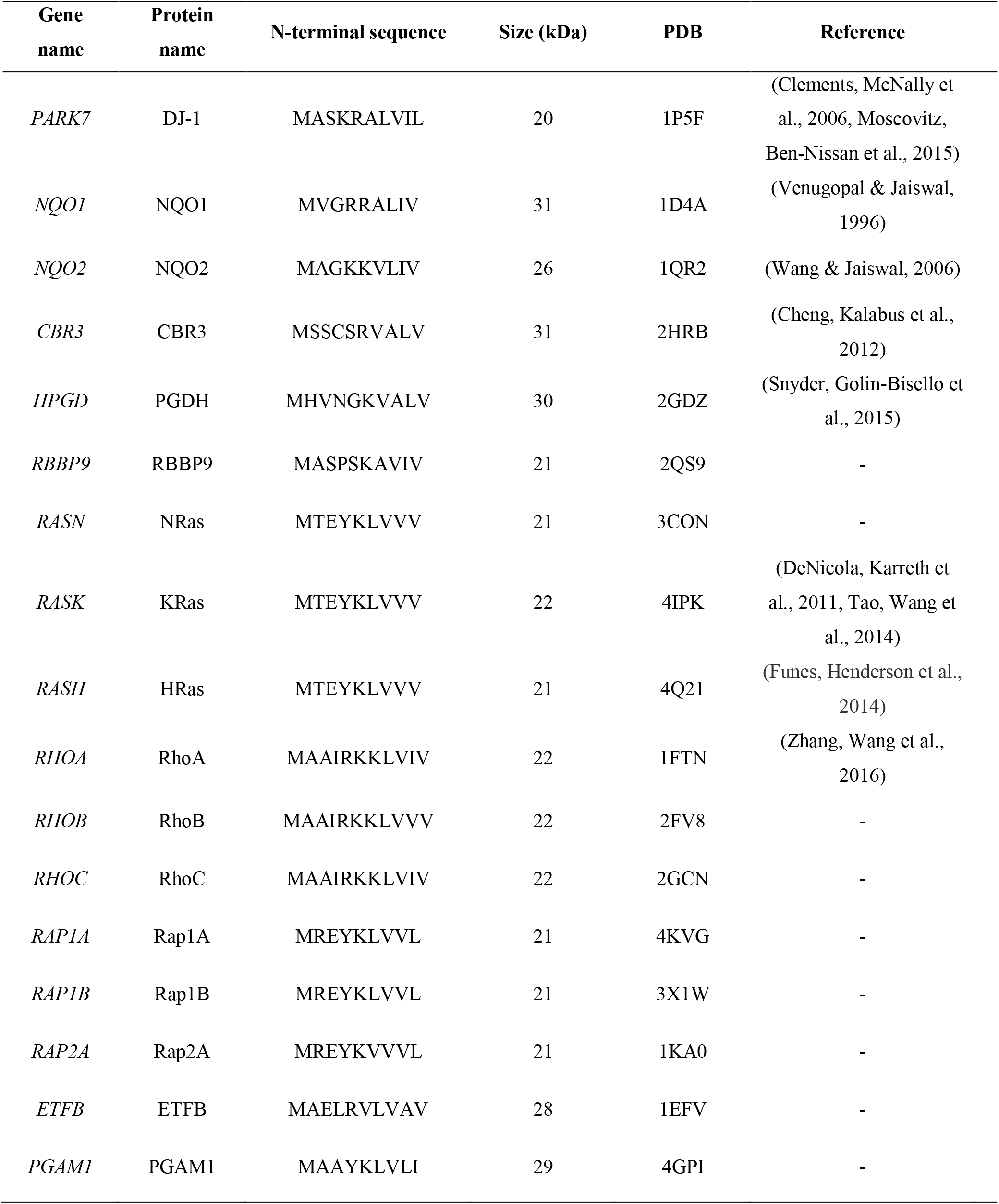
Shortlisted CCR candidates containing conserved N-terminal motif and a Rossman fold.

### CCRs inhibit 20S proteasome substrate degradation in vitro

To determine if any of these candidates are indeed capable of effecting 20S proteasome activity, we selected in addition to NQO1 and DJ-1, eight proteins for analysis: NQO2, CBR3, PGDH, RBBP9, NRas, KRas, HRas and RhoA. These proteins were expressed, purified and tested by *in vitro* degradation assays with purified mammalian 20S proteasomes and two different model substrates, α-syn (Tofaris, Layfield et al., 2001) and oxidized calmodulin (OxCaM) (Ferrington, Sun et al., 2001) (Figure 3A). MG132 was included as a control for proteasome inhibition. The majority of the CCRs successfully inhibited the degradation of α-syn, with the exception of RBBP9 and HRas. However, all of the candidates prevented the degradation of OxCaM. These results indicate that the CCRs are capable of inhibiting protein degradation by the 20S proteasome *in vitro*, with an element of substrate specificity demonstrated for HRas and RBBP9.

**Figure 3:**
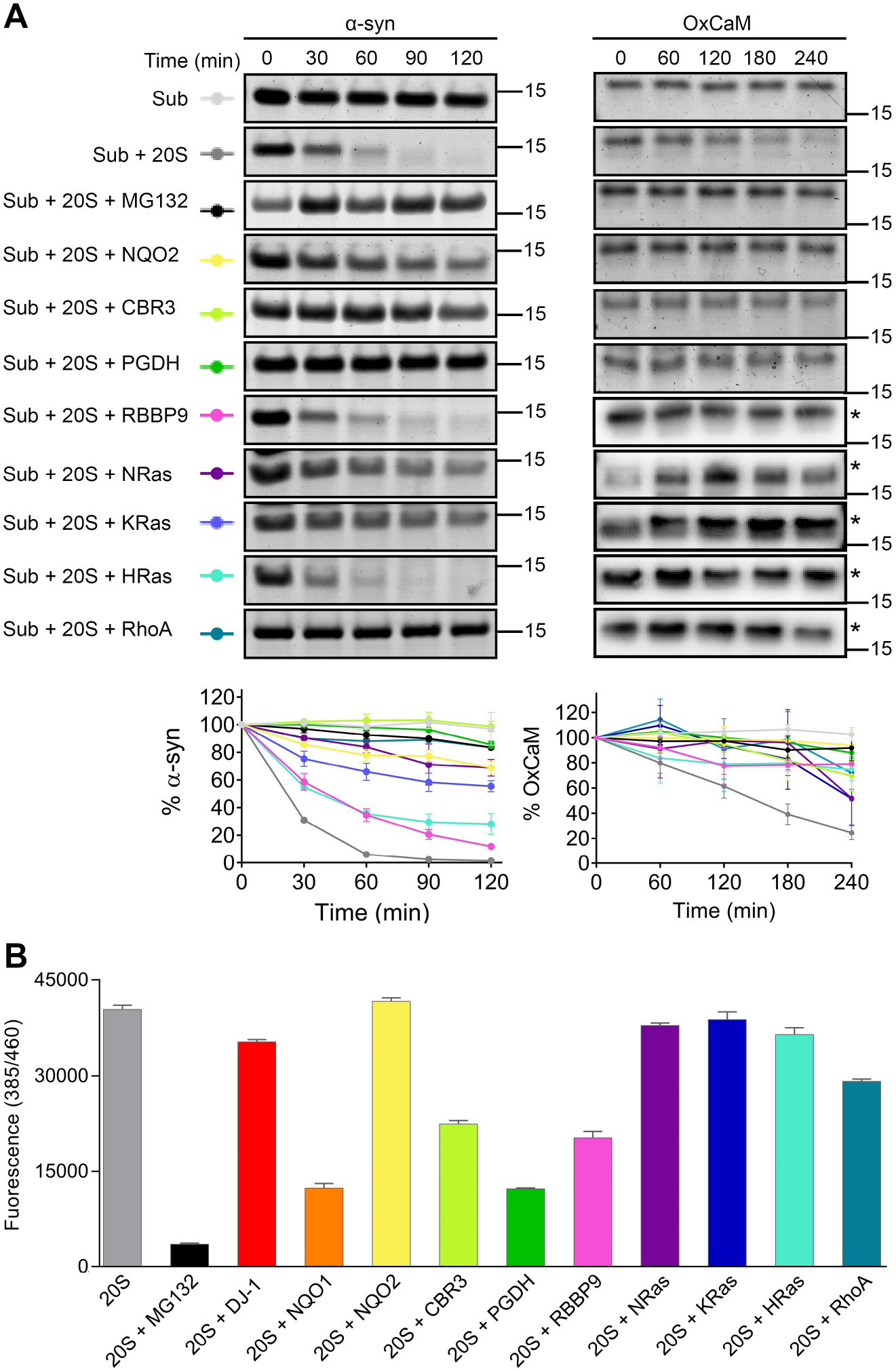
CCRs inhibit the degradation of partially folded proteins by the 20S proteasome. (A) *In vitro* degradation assays of each CCR with substrates (Sub) α-syn (left) or OxCaM (right). MG132 was included as a control for 20S proteasome inhibition. Panels display coomassie stained SDS-PAGE, unless labelled with an asterisk, which denotes immunoblots using anti-calmodulin antibody for those CCRs that are the same size as the substrate: RBBP9, NRas, KRas, HRas and RhoA. Quantification of three independent experiments is displayed, error bars represent S.E.M. (B) Peptidase activity of 20S proteasomes in the presence of CCRs or MG132 was monitored using the fluorogenic peptide substrate suc-LLVY-AMC. Error bars represent S.E.M. of three independent experiments.

To clarify whether the inhibition is a result of competitive inhibition i.e. the CCRs themselves are being degraded by the 20S proteasome in preference to the model substrates, each CCR was analyzed by *in vitro* degradation assay with 20S proteasome in the absence of a substrate (Supplementary Figure 2A). Quantification of the amount of CCR remaining over the course of the assay indicated that they themselves are stable, and are therefore not acting as competitive inhibitors (Supplementary Figure 2B).

We continued by examining whether the CCRs can affect the enzymatic activity of the 20S proteasome using a fluorogenic peptidase activity assay (Figure 3B). Compared with MG132, which drastically reduced the proteolytic activity of the 20S proteasome, several of the CCRs behaved like DJ-1 (Moscovitz et al., 2015), and showed an insignificant effect on peptide degradation. However, CBR3 and RBBP9 reduced the proteolytic activity by about half, while NQO1 and PGDH showed the most significant inhibition. This data suggests that while there are differences in the capacity of the CCRs to prevent peptide degradation, they are not able to completely block degradation, as was observed for MG132. This indicates that they do not deactivate the catalytic sites of the proteasome, but rather they act using another regulatory mechanism. Whether the CCRs prevent protein degradation by masking the entrance to the proteasome or by an allosteric mechanism remains to be determined.

To address whether the CCRs inhibitory capacity on the 20S proteasome is conserved across evolution, as was demonstrated for DJ-1 (Figure 2), we selected CBR3 as a representative CCR and analyzed its effect on the degradation of α-syn by yeast and archaeal 20S proteasomes (Supplementary Figure 3A-B). Human CBR3 (hCBR3) inhibited degradation by both proteasomes, indicating that like DJ-1, inhibition of the 20S proteasome by the CCR family is conserved across evolution. This observation further strengthen the possibility of conservation of function across CCRs from evolutionarily distant species.

### CCRs preferentially bind to the 20S proteasome over the 26S proteasome, to inhibit substrate degradation

The ability of the CCRs to inhibit protein degradation could be due to either direct interactions with the 20S proteasome, or sequestration of the substrate away from the proteasome by forming a stable complex with the regulator. Previous experiments with NQO1 and DJ-1 revealed that they did not interact with the substrate itself but rather physically bound to the 20S proteasome and inhibited its activity directly (Moscovitz et al., 2015, Moscovitz et al., 2012). We therefore utilized this native mass spectrometry approach and started by analyzing the interaction between the CCCRs and α-syn (Supplementary Figure 4). α-syn was incubated with each of the CCRs and their native mass spectra were analyzed. No larger complexes formed by the CCRs and α-syn were detected, suggesting that the inhibition of protein degradation does not occur by competitive inhibition via substrate sequestration.

The inhibition of protein degradation is therefore likely mediated by direct binding of the CCRs to the 20S proteasome. To test this, each of the CCRs were incubated with 20S proteasome, and tandem MS (MS/MS) was employed to detect binding. MS/MS involves three stages, beginning with the acquisition of a native MS spectrum of the intact protein complex in the protein mixture. This allows for the identification of the 20S proteasome in the high m/z range, as well as free CCR in the low m/z range. The peak series corresponding to the 20S proteasome complex is then isolated, allowing for specific selection of the 20S proteasome and its associated proteins, and not free CCR that remains unbound. The isolated complexes are subjected to high collision energies, leading to dissociation of any bound proteins as well as individual subunits of the 20S proteasome. These dissociated monomeric subunits and proteins can be detected in the low m/z range of the spectrum, and mass assignment allows for the identification of known 20S subunits, as well as CCRs that were bound to the 20S proteasome (Figure 4A). For each of the samples containing the CCRs, a unique series of peaks corresponding in size to the predicted molecular weight of the specific CCR were identified, that were not found in the spectrum for the 20S proteasome alone, alongside peak series corresponding to known 20S proteasome subunits (Figure 4B-K, Supplementary Table 1). This indicates that the CCRs bind directly to the 20S proteasome to regulate its function.

**Figure 4:**
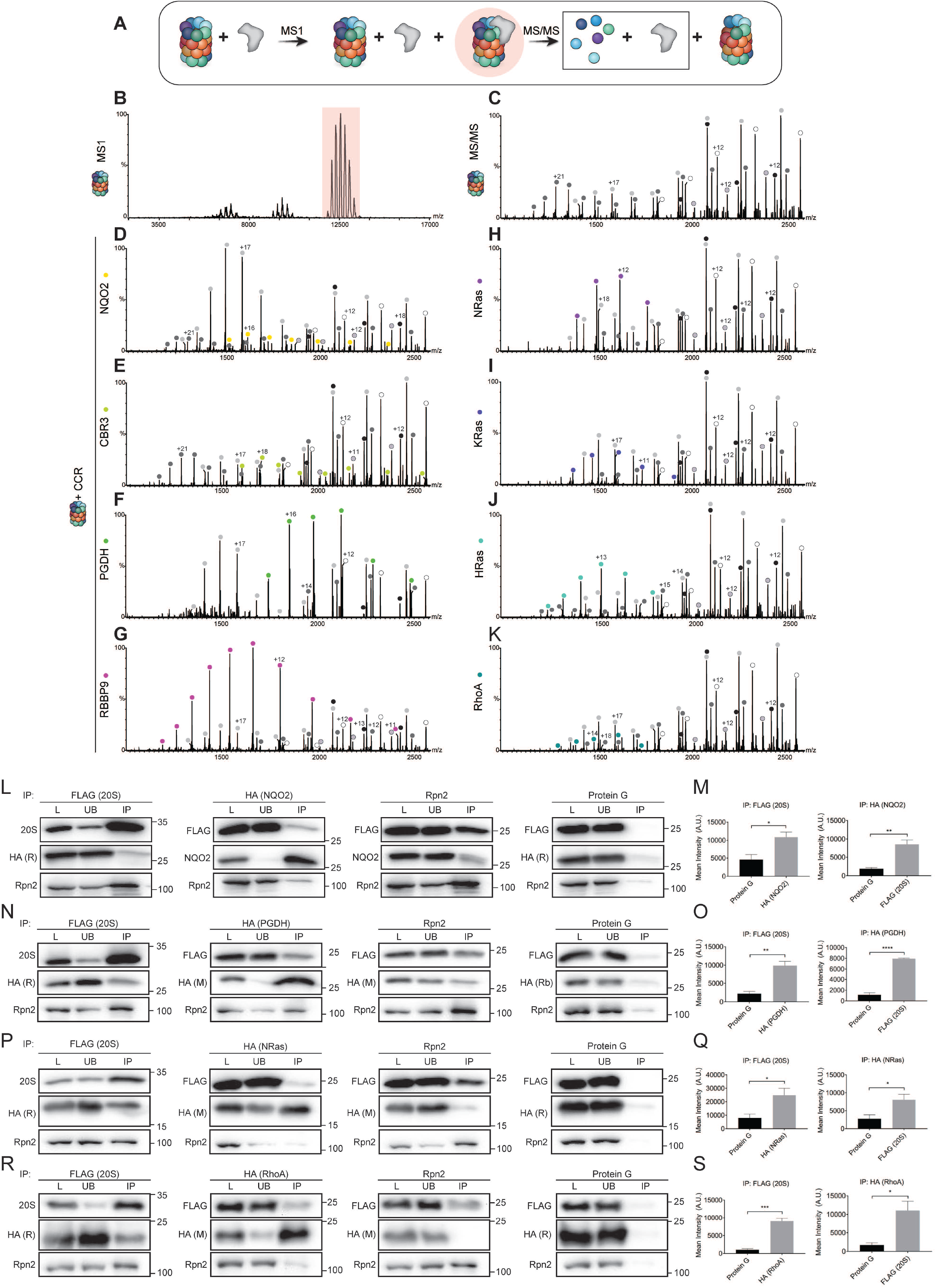
CCRs physically bind the 20S proteasome both *in vitro* and in cells. 20S proteasomes alone or in the presence of CCRs were analyzed by native mass spectrometry to determine the binding status of the CCRs to the 20S proteasome. (A) Schematic of native MS methodology, in which 20S proteasome are incubated with CCRs, leading to a mixed population of free 20S proteasomes, free CCR and CCR-bound 20S proteasome complexes. The complexes are isolated (highlighted in pink) and subjected to increased collision energy. This leads to the dissociation and detection of 20S proteasome subunits and bound CCRs (indicated by the box), leaving a stripped 20S proteasome lacking bound CCRs and several subunits. (B) Native MS1 spectrum of 20S proteasomes. Highlighted peaks were isolated and subjected to increased collision energy. (C) MS/MS spectrum of 20S proteasome, the charge series of individual dissociated 20S subunits were identified (white, grey, black balls). (D-K) 20S proteasomes were pre-incubated with CCRs, followed by MS/MS analysis to identify CCR binding. Unique charge series corresponding in size to the monomeric size of each of the CCRs was detected (colored balls), indicating that CCRs physically bind to the 20S proteasome. For predicted and measured masses of 20S proteasome subunits and CCRs, see Supplementary Table 1. For cellular experiments, HA-tagged (L-M) NQO2, (N-O) PGDH, (P-Q) NRas and (R-S) RhoA were overexpressed in HEK293T cells stably expressing FLAG-tagged B4 subunit of the 20S proteasome. For RhoA (R-S), cells were exposed to 100 µM DEM for 48 hours prior to collection and lysis. Lysates were subjected to IP using either anti-FLAG-affinity gel, anti-HA or anti-Rpn2 antibodies, or uncoupled Protein G beads as a control. Total starting lysate (L), unbound proteins (UB) and IP samples were analysed by western blot using anti-20S, anti-HA or anti-Rpn2 antibodies. Bands corresponding to HA (i.e. CCRs) in the FLAG (20S) IP and FLAG (20S) in the HA IP were quantified and compared with their Protein G equivalents. Each of the CCRs were significantly enriched in the FLAG (20S) IP demonstrating that the CCRs bind to the 20S proteasome. The reciprocal IP with anti-HA confirmed this interaction, with significant enrichment of the FLAG (20S) bands. Quantifications demonstrate the average of (M, O, S) four or (Q) five independent experiments. Band intensity measurements were subjected to Students t- test analysis, * p < 0.05, ** p < 0.01, *** p < 0.001, **** p < 0.0001. Error bars represent S.E.M.

To determine whether this binding occurs in cells, and is specific for the 20S proteasome, or if the CCRs can also bind to the 26S proteasome, immunoprecipitation (IP) experiments were performed using HEK293T cells stably expressing the 20S proteasome β_4_ subunit with a FLAG-tag on the C-terminus. Four representative CCRs (NQO2, PGDH, NRas and RhoA) were selected for the analysis. Each of these CCRs, containing a C-terminal HA-tag, were transiently overexpressed prior to lysis and IP to aid in the efficiency of the pull down. Whole cell lysates were IP’d with either anti-FLAG, anti-HA or anti-Rpn2 (a subunit of the 19S regulatory particle of the 26S proteasome) antibodies, to pull down the 20S proteasome, the CCRs or the 26S proteasome respectively. A control IP using uncoupled Protein G beads was performed in parallel to ascertain the background levels of the proteins binding to the beads during the IP. Bound proteins were then analyzed by western blotting with anti-PSMA1 (20S proteasome α_1_ subunit), anti-Rpn2 (19S subunit) and anti-CCR/HA antibodies (Figure 4L, N, P, R). Quantification of the levels of the CCRs being pulled down with the 20S proteasome in the FLAG IP revealed a significant increase in the amount of CCRs binding to the 20S proteasome, compared with the Protein G control (Figure 4M, O, Q, S – left panels). This interaction was confirmed in the reciprocal IP, where the amount of 20S proteasome being pulled down with the CCRs in the HA IP is significantly increased compared with the Protein G control (Figure 4M, O, Q, S – right panels). Of note, the interaction between RhoA with the 20S proteasome was detectable only after exposure of the cells to oxidative stress, likely due to the increased levels of 20S proteasome under these conditions, as discussed earlier. Binding of the CCRs to the 26S proteasome (Rpn2) only occurred at low levels in both the HA and Rpn2 IPs, indicating a preference for CCR binding to the 20S proteasome, or to singly capped 26S proteasomes. Taken together, these results establish that the CCRs specifically bind to the 20S proteasome in cells.

### CCRs stabilize the cellular levels of 20S proteasome substrates

We were motivated to examine whether our *in vitro* observations, revealing the CCRs ability to bind to the 20S proteasome and to prevent substrate degradation, are also relevant in a cellular context. To explore this, we overexpressed three representative CCRs (NQO2, CBR3 and PGDH), alongside GFP as a control, in HEK293T cells and measured the effect on known 20S proteasome substrates, α-syn and p53 (Asher et al., 2005) (Figure 5A-C). The cellular levels of both α-syn and p53 increased significantly when the CCRs were overexpressed compared with GFP. These data indicate that the 20S proteasome is being inhibited by the increased levels of CCRs in the cells, leading to reduced degradation of substrates and their consequent accumulation.

**Figure 5:**
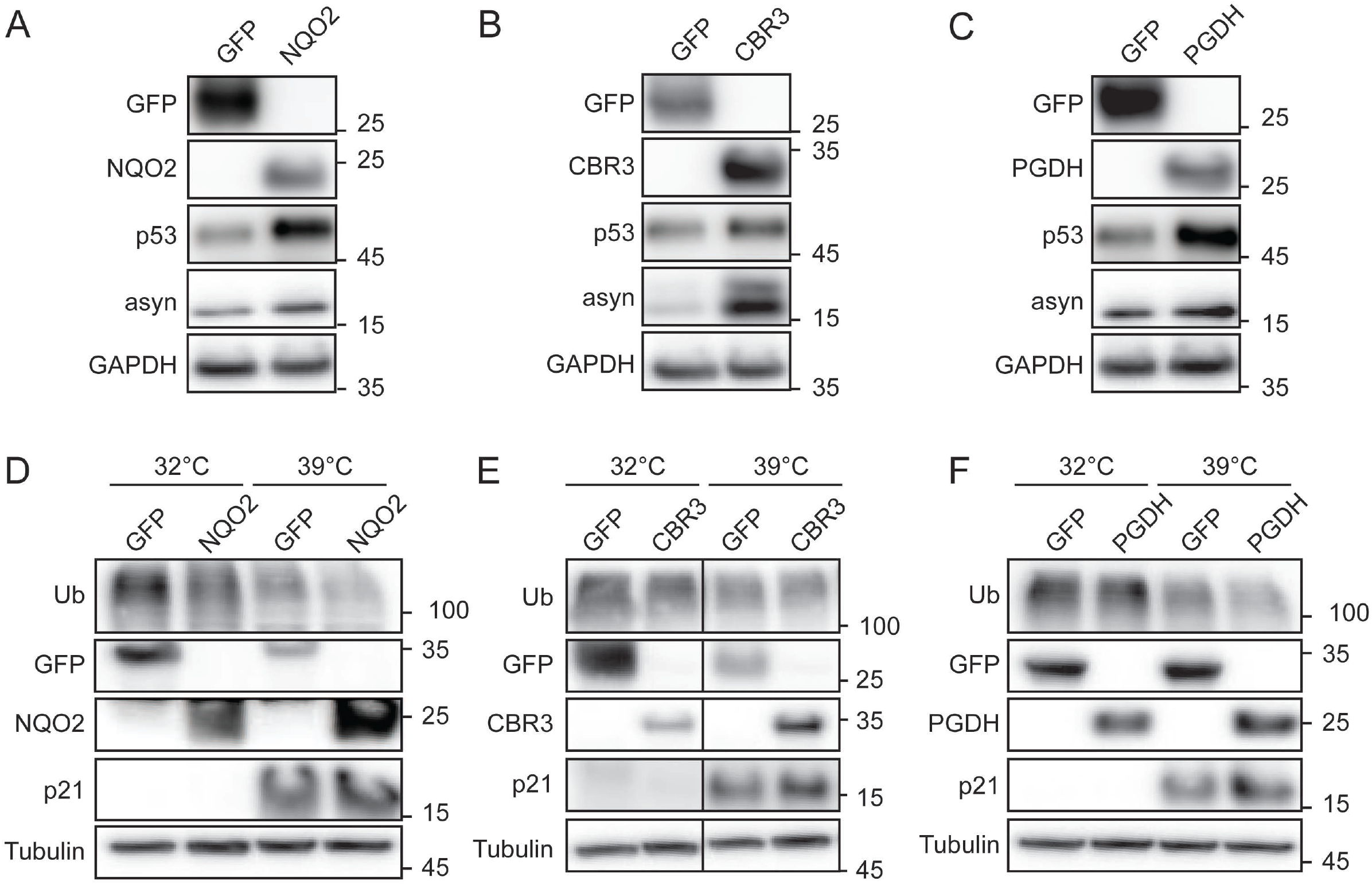
Overexpression of CCRs stabilizes the cellular levels of 20S proteasome substrates. Representative CCRs NQO2, CBR3 and PGDH were transiently overexpressed in HEK293T cells (A-C, respectively) and A31N cells (D-F, respectively). GFP was transfected in parallel as a control in all experiments. The levels of 20S proteasome substrates α-syn and p53 in HEK293T cells, and p21 in A31N cells were detected by western blot. p21 was only detected at the restrictive temperature of 39 °C in A31N cells due to inactivation of the ubiquitination cascade, as seen by the reduction in ubiquitin conjugates compared with 32 °C. In all cases, the levels of the substrates were enriched when the CCRs were overexpressed compared with GFP, indicating inhibition of 20S proteasome mediated degradation. Results are representative of three independent experiments.

To determine if the effect of CCR inhibition of the 20S proteasome is relevant to other known substrates, we followed the levels of a cell cycle regulator, p21, an IDP that has been previously shown to be a substrate of the 20S proteasome (Chen, Barton et al., 2007, Chen, Chi et al., 2004, Li, Amazit et al., 2007). However, analysis of this protein is complicated by two factors. First, evidence has shown that p21 can be ubiquitinated and degraded both by the 26S proteasome and the 20S proteasome (Bornstein, Bloom et al., 2003), making it challenging to differentiate by which route p21 is being degraded and therefore which degradation pathway is affected by any treatment or intervention. Second, the basal levels of p21 are low across most of the cell cycle, with the exception of enrichment at the G1/S checkpoint (Yoon et al., 2012), leading to difficulties in the detection of p21 at sufficient levels for analysis. Therefore, in order to follow changes in the levels of p21, we used the mouse fibroblast cell line, A31N-ts20 BALB/c, which contains a temperature sensitive mutant of the E1 ubiquitin activating enzyme (Salvat, Acquaviva et al., 2000). Incubation of these cells at the permissive temperate of 32 °C allows normal cell growth. Upon transfer of the cells to the restrictive temperature of 39 °C, the E1 enzyme is deactivated, and the ubiquitination cascade is subsequently affected. The reduction in ubiquitination levels allows for the accumulation of proteins such as p21, which is normally ubiquitinated and degraded. Under these conditions, degradation by the 26S proteasome is attenuated, while leaving the 20S proteasome degradation route unaffected. We transiently overexpressed the same three CCRs (NQO2, CBR3 and PGDH) in A31N-ts20 BALB/c cells, followed by 24 hours of growth at either 32 °C or 39 °C (Figure 5D-F). At the restrictive temperature, we clearly see a reduction in ubiquitin conjugate levels, and appearance of p21, due to the loss of the ubiquitination cascade. Compared with the GFP control, p21 levels are enriched when the CCRs are overexpressed. This stabilization of p21 levels is likely due to the specific inhibition of the 20S proteasome degradation pathway by the CCRs. Altogether, this data demonstrates that the CCRs influence the cellular levels of 20S proteasome substrates.

### CCRs and Nrf2 form a robust regulatory circuit during oxidative stress

Previous research demonstrated that NQO1 and DJ-1 form a feedback loop with the transcription factor Nrf2 in response to oxidative stress to regulate the activity of the 20S proteasome (Moscovitz et al., 2015). DJ-1 was shown to stabilize the levels of Nrf2 (Clements, McNally et al., 2006), which in turn induces the expression of NQO1 and subunits of the 20S proteasome (Moscovitz et al., 2015). In parallel, DJ-1 and NQO1 both inhibit the 20S proteasome, thus providing a delicate balance between increased 20S proteasome levels during oxidative stress, and appropriate control of 20S proteasome degradation (Moscovitz et al., 2015). Many of the newly discovered CCRs have been shown to be linked to Nrf2 either as downstream transcriptional targets or upstream factors that influence its stability (Cheng, Kalabus et al., 2012, DeNicola, Karreth et al., 2011, Funes, Henderson et al., 2014, Snyder, Golin-Bisello et al., 2015, Tao, Wang et al., 2014, Wang & Jaiswal, 2006, Zhang, Wang et al., 2016). To determine the status of each CCR during the Nrf2 response to oxidative stress, we down regulated the levels of Nrf2 followed by induction of oxidative stress (Figure 6A-B). Quantification of the levels of the CCRs lead us to classify them into three groups, depending on their response to oxidative stress and Nrf2 silencing (Figure 6C-E). The first group are the inducible Nrf2 responders: NQO2 and CBR3 responded in the same manner as NQO1, with increasing protein levels over the time-course of oxidative stress, which were reduced when Nrf2 was silenced (Figure 6C). The second group is the basal Nrf2 responders, comprising only of NRas, which did not show a significant increase during oxidative stress, however when Nrf2 was silenced its levels were significantly reduced (Figure 6D). This observation is supported by previous studies showing that Nrf2 can activate the transcription of certain genes under basal conditions, as opposed to under oxidative stress (Malhotra, Portales-Casamar et al., 2010). This is the first reported data revealing transcriptional regulation of NRas by Nrf2, hinting towards a potentially important influence of Nrf2 on the Ras family signaling pathways.

**Figure 6:**
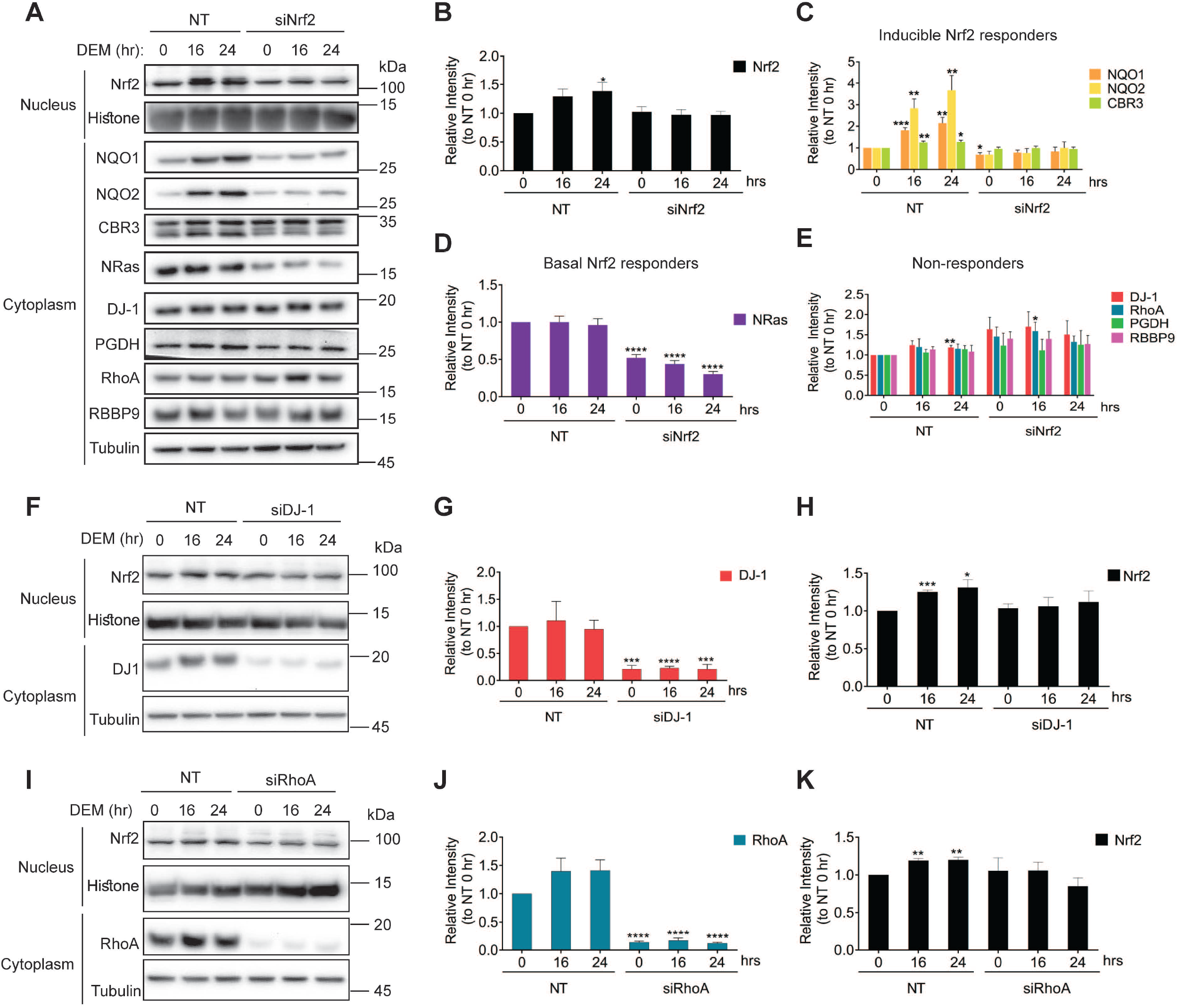
CCRs are involved in the oxidative stress response via the Nrf2 pathway. (A) MCF10A cells were transiently transfected with either non-targeting siRNA (NT) or siRNA against Nrf2 (NFE2L2). 48 hours post transfection, cells were exposed to 150 µM DEM to induce oxidative stress for 0, 16 or 24 hours. Cells were collected, fractionated into cytoplasmic and nuclear fractions, and analyzed by western blot to assess the levels of Nrf2 and the CCRs. (B-E) Changes in the levels of Nrf2 and the CCRs relative to NT at 0 hours were quantified from 4 independent experiments. The CCRs were categorized into three groups based on their response to Nrf2 silencing; (C) inducible Nrf2 responders (NQO1, NQO2, CBR3), (D) basal Nrf2 responders (NRas) and (E) non-responders (DJ-1, RhoA, PGDH and RBBP9). The CCRs DJ-1 and RhoA from the non-responder group were transiently silenced in MCF10A cells for 48 hours, followed by exposure to DEM for 0, 16 or 24 hours. Cells were collected, fractionated into cytoplasmic and nuclear fractions, and analyzed by western blot to assess the levels of Nrf2 during (F) DJ-1 and (I) RhoA silencing. Changes in the levels of (G) DJ-1 and (H) Nrf2 during DJ-1 silencing, and (J) RhoA and (K) Nrf2 during RhoA silencing, demonstrates that these two CCRs influence the translocation of Nrf2 to the nucleus during oxidative stress. Altogether, this data indicates that the CCRs organize into a regulatory loop with Nrf2, providing tight control over the 20S proteasome during oxidative stress. Quantification relative to NT at 0 hours was performed from three independent experiments. Band intensity measurements were subjected to Students t-test analysis, * p < 0.05, ** p < 0.01, *** p < 0.001, **** p < 0.0001. Error bars represent S.E.M.

The CCRs (along with DJ-1) that did not increase during oxidative stress, or decrease when Nrf2 was silenced, and are thus classified as non-responders, were PGDH, RBBP9 and RhoA (Figure 6E). Interestingly, the presence of DJ-1 and RhoA in this group adds additional significance to the tissue wide expression data (Supplementary Figure 1), which demonstrated that these two CCRs show the highest and most widespread protein expression levels of all the CCRs analyzed. Given that DJ-1 was shown to stabilize Nrf2 (Moscovitz et al., 2015), there could be a correlation between widespread protein expression and Nrf2 stabilization. To determine if RhoA can stabilize the levels of Nrf2 in the same manner as DJ-1, we silenced these CCRs and monitored the levels of Nrf2 in the nucleus during oxidative stress. As expected, silencing of DJ- 1 prevented an increase in Nrf2 in nucleus during oxidative stress (Figure 6F-H), in accordance with our previous results (Moscovitz et al., 2015). Reducing the levels of RhoA also significantly weakened the response of Nrf2 to oxidative stress, preventing nuclear accumulation of Nrf2 (Figure 6I-K). These results indicate that DJ-1 and RhoA act upstream of Nrf2 to stabilize its levels and ensure appropriate nuclear accumulation during oxidative, enabling the oxidative stress response to occur.

## Discussion

This study describes the discovery of a novel family of 20S proteasome regulators, the CCRs. We have demonstrated that a robust link exists between Nrf2 and the CCRs, which specifically regulate the 20S proteasome by physically binding, and attenuating, substrate degradation. Nrf2 controls the expression of the some of these CCRs, while Nrf2 itself is stabilized by at least two other members of the CCR family. Together, this creates a feed forward loop of regulation (Mangan & Alon, 2003), within which the 20S proteasome is upregulated by Nrf2 and simultaneously inhibited by the CCRs (Figure 7). This enables a pulse of activity and rapid shutdown during the oxidative stress response, allowing degradation of damaged proteins while protecting the levels of important regulatory proteins and preventing proteasome clogging, to ensure recovery after the oxidative insult has subsided.

**Figure 7:**
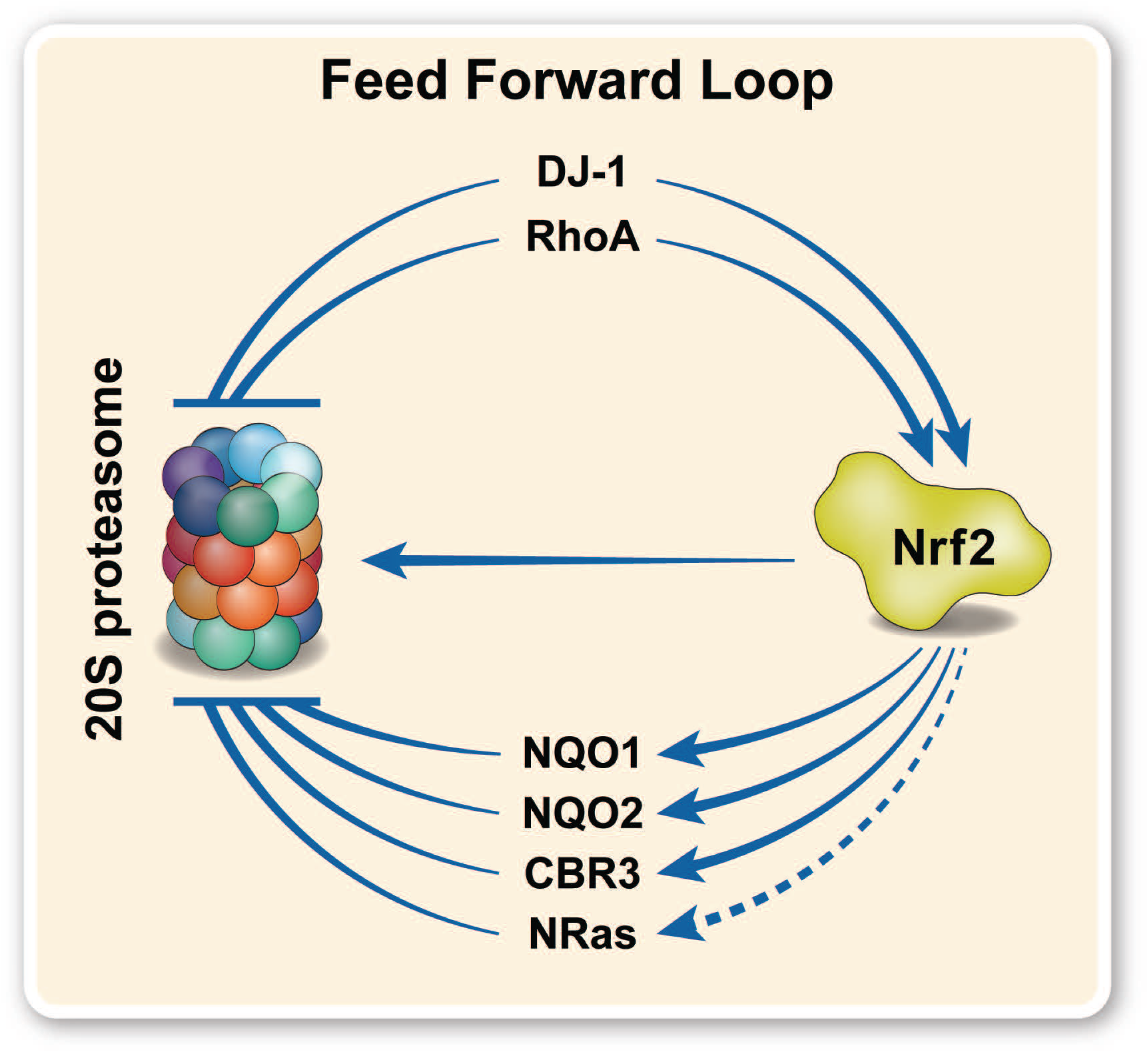
CCRs organize into a feed forward regulatory loop with Nrf2 and the 20S proteasome. During oxidative stress, Nrf2 translocates to the nucleus to initiate the oxidative stress response by transcription of target genes, including subunits of the 20S proteasome and the CCRs NQO1, NQO2 and CBR3. NRas is basally transcribed by Nrf2 (indicated by dashed line). DJ-1, RhoA and PGDH act upstream of Nrf2 to stabilize its levels. All the CCRs concurrently inhibit the 20S proteasome, preventing the degradation of its substrates and dampening the effect of the increased 20S proteasome levels during oxidative stress, protecting the cell from damaging imbalances and proteasome clogging, and ensuring a rapid recovery.

The members of the CCR family described in this study are greatly varied in their alternative functions beyond their moonlighting roles as 20S proteasome inhibitors. Multiple roles for DJ-1 have been described, such as DNA repair (Richarme, Liu et al., 2017) and methylglyoxylase activity (Toyoda, Erkut et al., 2014). NQO1 and NQO2 are both members of the quinone reductase family (Dinkova-Kostova & Talalay, 2010, Wang & Jaiswal, 2006), while other CCRs are involved in various metabolic pathways, such as carbonyl reduction (CBR3) (Schaupp, White et al., 2015), prostaglandin inactivation (PGDH) (Tai, Cho et al., 2006) and serine hydrolysis (RBBP9) (Vorobiev, Huang et al., 2012). In addition, multiple members of the Ras superfamily, which are small GTPases that are involved in cell proliferation, morphology, signaling, transport etc (Goitre, Trapani et al., 2014), are represented in the CCRs. In addition to those CCRs that were analyzed in this study, there are additional candidates that still need to be explored (Table 1). These include more members of the Ras superfamily, electron transfer flavoprotein B (ETFB), the β subunit of the electron transfer flavoprotein complex in the mitochondria, known to generate free radicals and therefore contribute to oxidative stress (Rodrigues & Gomes, 2012), and phosphoglycerate mutase 1 (PGAM1), an enzyme in the glycolytic pathway (Chaneton & Gottlieb, 2012). The varying roles of the CCRs, along with the ability to inhibit the 20S proteasome, suggest that the dual functionality of these proteins may be important for focused inhibition of the 20S proteasome in association with specific metabolic or signaling pathways, in a spatiotemporal manner. In support of this hypothesis, research has shown that RBBP9 is located in the extranuclear compartment (Shields, Niessen et al., 2010), CBR3 and NQO2 are cytoplasmic (Miura, Nishinaka et al., 2008, Wang & Jaiswal, 2006), RhoA is associated with the cytoskeleton and NRas, KRas and HRas shuttle between the cytoplasmic membrane and the cytosol depending on their nucleotide state (Goitre et al., 2014). It has been shown that the free, constitutive 20S proteasome also localizes to different cellular compartments (Ben-Nissan & Sharon, 2014, Breusing & Grune, 2008, Fabre et al., 2013), therefore the presence of location-specific regulators may be vital to ensure that 20S proteasomal degradation occurs at an appropriate pace and on the necessary substrates in each location in the cell.

Many of the CCRs are also associated with mutation or expression driven pathologies, such as neurodegenerative disease and various cancers. For example, missense mutations in DJ-1 are known to cause early-onset Parkinson’s disease (Blackinton, Ahmad et al., 2005, Cookson, 2012, Kahle, Waak et al., 2009), and intriguingly, one of these mutants was shown to be able to inhibit the 20S proteasome more effectively than wild type DJ-1 (Ben-Nissan, Chotiner et al., 2016). Mutations in NQO1 have been correlated with an increased risk of Alzheimer’s disease (Bian, Zhao et al., 2008, Tsvetkov, Adamovich et al., 2011), while changes in PGDH expression have been linked to tumor growth in multiple cancers (Tai, Tong et al., 2007, Wolf, O’Kelly et al., 2006), as well as late-stage neurodegenerative disease (Miyagishi, Kosuge et al., 2017). Preliminary links have been found between changes in RBBP9 levels and ALS progression (Lilo, Wald-Altman et al., 2013) as well as pancreatic cancer (Shields et al., 2010, Vorobiev et al., 2012). RhoA overexpression is associated with cancer progression (Zhang et al., 2016), and the Ras family of proteins are famously associated with numerous oncogenetic mutations, leading to constitutively activated Ras proteins that drive the development and growth of multiple cancers (Hobbs, Der et al., 2016). Moreover, many cancers and neurodegenerative diseases are also associated with oxidative stress (Kim, Kim et al., 2015, Reuter, Gupta et al., 2010), and as described here, most of the CCRs analyzed are involved in the oxidative stress response. It is therefore tempting to speculate that given the dual roles of these moonlighting proteins, mutations or changes in their expression may also affect their ability to inhibit the 20S proteasome and participate in the oxidative stress response, thus leading to imbalances in the levels of 20S proteasome substrates, compounding the negative effects on the cell.

The discovery of the CCR family was based on the presence of a Rossman fold and the conserved N-terminal sequence motif found across NQO1 and DJ-1 homologues (Figure 3). Considering that Rossman folds are one of the most common structural motifs found in proteins (Hanugoklu, 2015, Laurino, Toth-Petroczy et al., 2016), it is likely that many more proteins exist that share these features, but whose structures have yet to be elucidated. In addition, the Rossman fold is one of the most ancient protein folds, found even in ancient organisms such as Archaea (Caetano-Anolles, Kim et al., 2007, Ma, Chen et al., 2008). The conservation of function of the CCRs in regulating the 20S proteasome from Archaea (Figure 2, Supplementary Figure 3) indicates that this regulatory interaction may have evolved early in evolution, possibly even before the evolution of the 26S proteasome pathway, demonstrating the importance of modulating the degradation of proteins by the 20S proteasome. Therefore, the CCR family may actually contain many more members than those described here, making the regulatory network of the 20S proteasome far more wide reaching than currently thought. Future research into the CCRs analyzed in this study, and the potential discovery of more members of this family, will be critical for mapping the global network of proteasome regulators, as well as for elucidating the exact mechanistic details of 20S proteasome regulation.

The clinical success of proteasome inhibitors, such as Bortezomib for the treatment of multiple myeloma and mantle cell lymphoma, is testament to the potential of the proteasome pathways as drug targets (Utecht & Kolesar, 2008). However, the inability of these proteasome inhibitors to distinguish between the 26S and 20S proteasomes, which as described here display different functions and are regulated by different mechanisms, has been a drawback of these therapies, leading to unwanted and deleterious side effects (Frankland-Searby & Bhaumik, 2012). As such, exploiting this novel 20S proteasome specific regulation, possibly via a conserved mechanism shared by the CCRs, could open avenues for the development of novel 20S proteasome specific inhibitors to treat diseases of proteasome dysfunction.

## Acknowledgements

We thank Sarel Fleishmann and Gideon Schreiber for helpful discussions. We thank Yardena Samuels for generously providing NRas antibodies and the pCDF1 plasmid, and for helpful discussions. We thank Yossi Shaul for generously providing MCF10A cells. We thank Ron Rotkopf and Maria Fuzesi-Levi for helpful insights and discussions. We are also grateful for the support of a Starting Grant from the European Research Council (ERC) (Horizon 2020)/ERC Grant Agreement No. 636752. M.A.O is supported by a Deans Fellowship from the Faculty of Life Sciences, a Senior Postdoctoral Fellowship from the Koshland Fund, Weizmann Institute of Science, and a Postdoctoral Fellowship from the Israel Cancer Research Fund. M.S. is an incumbent of the Aharon and Ephraim Katzir Memorial Professorial Chair.

## Author Contributions

Conceptualization, M.A.O. and M.S.; Methodology, M.A.O., F.K.D., G.A., I.F., G.B.N., M.S.; Formal Analysis, M.A.O., F.K.D., G.A., I.F., D.H., S.B.D., G.B.N.; Investigation, M.A.O., F.K.D., G.A., I.F., M.T., D.H., S.B.D, G.B.N.; Resources, S.B.D., M.S.; Writing – Original Draft, M.A.O. and M.S.; Writing – Review & Editing, M.A.O., F.K.D., G.A., I.F., S.B.D., G.B.N., M.S.; Visualization, M.A.O.; Supervision, G.B.N., M.S.; Funding Acquisition, M.A.O., M.S.

## Competing Interests

The authors declare no competing interests.

## Methods

### Cell lines

HEK293T cells stably expressing the β4 subunit of the proteasome tagged with a C-terminal FLAG tag were obtained from Chaim Kahana (Weizmann Institute of Science). Wild type HEK293T cells were obtained from Eitan Reuveny (Weizmann Institute of Science). The mouse fibroblast cell line A31N-ts20 BALB/c and human mammary epithelial MCF10A cells were obtained from Yosef Shaul (Weizmann Institute of Science).

HEK293T and A31N-ts20 BALB/c cells were maintained in Dulbecco’s Modified Eagle’s Medium (Sigma) supplemented with 10% fetal bovine serum (Gibco), 100units/ml penicillin-100µg/ml streptomycin (Biological Industries), 0.1mM sodium pyruvate (Biological Industries), MEM-Eagle non-essential amino acids (Biological Industries) and MycoZap Prophylactic (Lonza) according to the manufacturer’s instructions. β4-FLAG HEK293T cells were additionally supplemented with 1mg/ml Puromycin. MCF10A cells were maintained in Dulbecco’s Modified Eagle’s Medium/Nutrient Mixture F-12 Ham (Sigma) supplemented with 5% Donor Horse Serum (Gibco), 20ng/ml EGF (Sigma), 10µg/ml insulin (Sigma), 0.5µg/ml hydrocortisone (Sigma), 100ng/ml cholera toxin (Sigma), 100units/ml penicillin-100µg/ml streptomycin (Biological Industries) and 2mM L-Glutamine (Biological Industries). Cells were grown in a humidified incubator at 37°C (HEK293T, MCF10A) or 32°C (A31N-ts20 BALB/c) with a 5% CO_2_ controlled atmosphere.

### Microbe strains

The DH5α strain of *E.coli* was used for all plasmid cloning experiments. The BL21(DE3) strain of *E.coli* was used for all recombinant protein expression experiments.

### Organisms as source for materials used in experiments

The *S. cerevisiae* strain RJD1144, generated in the lab of Raymond Deshaies, Caltech, California, USA, which contains a chromosomally FLAG-tagged β4 (PRE1) subunit was used for the purification of yeast 20S proteasome (Verma, Chen et al., 2000).

The *R. norvegicus* strain RCS was used as a source of liver for the purification of the mammalian 20S proteasome.

### Bioinformatic analyses

Sequences of DJ-1 and NQO1 homologues from multiple species were acquired from UniProtKB. Multiple sequence alignments (MSAs) were performed using Clustal Omega, figures were generated using ESPript (espript.ibcp.fr) and Weblogo (weblogo.berkeley.edu/logo.cgi). All CATH-annotated Rossman fold containing proteins were imported from the Protein Data Bank (www.rcsb.org), and the presence of the N-terminal motif (MX(1,4)[KR](1,2)[AVIL]4) discovered in the MSAs was identified using FuzzPro (EMBOSS (Rice, Longden et al., 2000)). Estimated protein expression levels across various human tissues were taken from the Human Integrated Protein Expression Database (HIPED) (Fishilevich, Zimmerman et al., 2016).

### Plasmids and cloning

The following plasmids were acquired from Addgene: pT7-7 α-syn WT (#36046), pNIC28-Bsa4 CBR3 (#38800), pET28-MHL NRASA (#25256, containing truncated NRas amino acids 1-172), pET28-MHL KRASB (#25153, containing truncated KRas amino acids 1-169, and Q61H mutation), pGEX2T-HRas (#55653), pNIC28-Bsa4 RhoA (#73231, containing truncated RhoA amino acids 1-184). pGEM-T NQO2 (HG14634-G) was acquired from Sino Biological. Modifications made to the plasmids are as follows: pGEX2T-HRas was used as a template to amplify truncated HRas (amino acids 1-172) using the primer pair: forward 5’AACCTGTACTTCCAGGGTACCATGACAGAATACAAGCTTGTG and reverse 5’CTCGAGTGCGGCCGCAAGCTTTCAGTTCAGTTTCCGCAAT. The amplified product was inserted into the pET28 plasmid using Infusion, introducing an N-terminal 6xHis tag and TEV cleavage site. pET28-MHL KRASB was used as a template to reverse the Q61H mutation, using the primer pair: forward 5’ CAGCAGGTCAGGAGGAGTACA and reverse 5’ TGTCGAGAATATCCAAGAGAC. pGEM-T NQO2 was used as a template to amplify NQO2 using the primer pair: forward 5’ AACCTGTACTTCCAGGGTACCATGGCAGGTAAGAAAGTACTCA and reverse 5’ CTCGAGTGCGGCCGCAAGCTTTCATTGCCCGAAGTGC. The amplified product was inserted into the pET28 plasmid using Infusion, introducing an N-terminal 6xHis tag and TEV cleavage site.

Total RNA was extracted from HEK293T cells using NucleoZOL (Macherey-Nagel) according to the manufacturer’s instructions. cDNA was synthesized using Protoscript II Reverse Transcriptase (NEB) and d(T)_20_ oligonucleotide according to the manufacturer’s instructions. RBBP9 and PGDH were amplified from the cDNA using the primer pairs: RBBP9 forward 5’ AACCTGTACTTCCAGGGTACCATGGCTTCTCCTAGCAAGGCA and reverse 5’ CTCGAGTGCGGCCGCAAGCTTCTATGCTGGTACTTTCAGCAA, PGDH forward 5’ AACCTGTACTTCCAGGGTACCATGCACGTGAACGGCAAAGTG and reverse 5’ CTCGAGTGCGGCCGCAAGCTTTCATTGGGTTTTTGCTTGAAATGGA. The amplified products were cloned into pET28 to introduce a C-terminal 6xHis tag.

The pCDF1 expression plasmid, used for mammalian cell transfection, was loaded with the following inserts amplified with their respective primers pairs; NQO2: forward 5’ GGGCGCGCCGGATCCATGGCAGGTAAGAAAGTACTCATTG and reverse 5’CGCGGCCGCGAATTCTCATTGCCCGAAGTGCCAG, NQO2-HA: forward as for NQO2 and reverse 5’ TCGCGGCCGCGAATTCTCACGCATAGTCAGGAACATCGTATGGGTAACCTCCTTGCCCGAAGTGCCAGTGGGC, CBR3: forward 5’ GGGCGCGCCGGATCCATGTCGTCCTGCAGCCGC and reverse 5’ CGCGGCCGCGAATTCTCACCAGTTTTGCACAACTTTG, PGDH: forward 5’ GGGCGCGCCGGATCCATGCACGTGAACGGCAAAGTG and reverse 5’ CGCGGCCGCGAATTCTCATTGGGTTTTTGCTTGAAATGG, PGDH-HA: forward as for PGDH and reverse 5’TCGCGGCCGCGAATTCTCACGCATAGTCAGGAACATCGTATGGGTAACCTCCTTGGGTTTTTGCTTGAAATGGAGTTG, NRas-HA: forward 5’ GAGCCCGGGCGCGCCGGATCCGCCACCATGACTGAGTACAAACTGGTG and reverse 5’ TCGCGGCCGCGAATTCTCACGCATAGTCAGGAACATCGTATGGGTAACCTCCGTTGAGTTTTTTCATTCGGTACTGG, RhoA-HA: forward 5’ CGGGCGCGCCGGATCCATGGCTGCCATCCGGAAG and reverse 5’ TCGCGGCCGCGAATTCTCACGCATAGTCAGGAACATCGTATGGGTAACCTCCCCCACGTCTAGCTTGCAGAGC. Empty pCDF1 and pCFD1-GFP were obtained from Yardena Samuels (Weizmann Institute of Science).

### Protein expression and purification

### Purification of substrates α-syn and OxCaM

BL21(DE3) were transformed with pT7-7 α -syn WT. Cells were grown in LB medium supplemented with 100 ug/ml ampicillin at 37 °C until they reached OD_600_ 0.8. Protein expression was induced by the addition of 1 mM isopropyl-β-D-thiogalactoside (IPTG) for 3 hours at 37 °C, after which the cells were moved to 16 °C overnight. Cells were harvested by centrifugation at 5000 *g* for 10 minutes, and resuspended in 50 mM Tris-HCl pH 8, 50 mM KCl, 5 mM MgAcetate, 10 mM EDTA pH 8. Cells were lysed in a French Press and centrifuged at 12,000 *g* for 20 minutes to remove cellular debris. The supernatant was boiled for 15 minutes, followed by centrifugation at 12,000 *g* for 20 minutes to remove insoluble proteins. The supernatant was collected and passed over a Q anion exchange column pre-equilibrated with 20mM Tris pH 8. After loading, the column was sequentially washed with 13%, 55%, 70% and 100% 20 mM Tris pH 8, 1M NaCl to elute the protein. Fractions containing α -syn were pooled and concentrated using a 3-kDa Amicon Ultra column (Millipore). Concentrated α -syn was loaded onto a gel filtration column (Superdex 200, 10/300 GL, GE Healthcare), pre-equilibrated with 50 mM Tris pH 8, 150 mM NaCl. Fractions containing α -syn were collected, concentrated, frozen in liquid nitrogen and stored at −80°C.

Calmodulin was purchased from Sigma as a lyophilized powder (P1431) and oxidized to produce OxCaM as previously described (Ferrington et al., 2001) with the following modification: instead of 50mM HOMOPIPES pH 5, 50mM HEPES pH 7.5 was used.

### Purification of human DJ1 and yeast DJ-1 (Hsp32)

DJ-1 and Hsp32 were expressed and purified as previously described (Moscovitz et al., 2015). BL21(DE3) were transformed with pET-15b-hDJ-1 or pET28-Hsp32. Cells were grown in LB medium supplemented with 100 ug/ml ampicillin or 50 µg/ml kanamycin respectively, at 37 °C until they reached OD_600_ 0.45. Protein expression was induced by the addition of 0.4 mM IPTG for 2.5 hours. Cells were harvested by centrifugation at 5000 *g* for 10 minutes, and resuspended in 50 ml of 50 mM Tris-HCl pH 7.4, 2 mM EDTA, 1 mM DTT, 1 mM PMSF and a protease inhibitor cocktail (Complete, Roche). Cells were lysed in a French Press, centrifuged for 10 min at 5000 *g* and the lysate was passed through a Source-15Q anion exchange 55 ml column (GE Healthcare) pre-equilibrated with 50 mM Tris-HCl pH 7.4, 1 mM DTT. After lysate loading, proteins were eluted with 200 ml of 50 mM Tris-HCl pH 7.4, 1 mM DTT. 50ml fractions were collected and DJ-1/Hsp32-containing fractions (eluted after 150–200 ml) were concentrated using a 3-kDa Amicon Ultra column (Millipore). Concentrated DJ-1/Hsp32 was loaded onto a gel filtration column (Superdex 200, 10/300 GL, GE Healthcare), pre-equilibrated with 50 mM Tris-HCl pH 7.4, 300 mM NaCl and 1 mM DTT. DJ-1/Hsp32-containing fractions were combined, concentrated, frozen in liquid nitrogen and stored at 80°C.

### Purification of archaea DJ-1

The BL21 (DE3) was transformed with pET28-taDJ-1. Cells were grown at 37°C to an OD_600_ of 0.5 in 100 ml LB medium supplemented with 50 µg/ml kanamycin. Protein expression was induced by the addition of 0.5 mM IPTG for 7 h at 37°C and then the cells were moved to 16°C for overnight protein expression. Cells were harvested by centrifugation for 20 min at 5000 *g* and resuspended in 50 mM Tris-HCl pH 7.5, 50 mM NaCl, 20 mM Imidazole, 250 U Benzonase (Millipore) 1 mM PMSF. Cells were lysed by sonication and the lysate was centrifuged for 30 min at 40,000 *g*. The supernatant was loaded onto a HisTrap FF 5 ml column (GE Healthcare) pre-equilibrated with 50 mM Tris-HCl, 50 mM NaCl, 20 mM Imidazole. After lysate loading, protein was eluted with 0-100% gradient elution buffer (50 mM Tris-HCl pH 7.5, 300 mM NaCl, 500 mM Imidazole). taDJ-1 containing fractions were pooled and dialyzed with TEV protease against 50 mM Tris pH 7.4, 1 mM EDTA and 2 mM DTT. Following the overnight TEV cleavage, the taDJ-1 was loaded on HisTrap FF 5ml and flow through fraction was collected, concentrated, frozen in liquid nitrogen and stored at −80°C.

### Purification of NQO2

NQO1 and NQO2 were expressed and purified as described previously with minor modifications (Moscovitz et al., 2012). BL21(DE3) were transformed with the pET28-NQO1 or pET28-NQO2. Cells were grown in LB medium supplemented with 50 µg/ml kanamycin at 37 °C until they reached OD_600_ 0.45. Protein expression was induced by the addition of 0.4 mM IPTG for 2.5 hours. Cells were harvested by centrifugation and lysed by sonication in 50 mM Tris-HCl (pH 7.5), 150 mM NaCl, 1 mM PMSF. Cellular debris was removed by sonication, and the supernatant was passed over a HisTrap FF column (GE Healthcare). His-TEV-NQO1 or His-TEV-NQO2 were eluted over a gradient up to 400 mM Imidazole. Fractions containing the proteins were pooled and TEV protease was added. The sample was incubated at room temperature for 4 hours, followed by overnight dialysis against 50 mM Tris-HCl (pH 7.5), 150 mM NaCl. The dialyzed protein was passed over a HisTrap FF column to remove the TEV protease and any uncleaved protein. The flowthough was collected and concentrated using a 10 kDa Amicon Ultra column (Millipore), and loaded onto a gel filtration column (Superdex 200, 10/300 GL, GE Healthcare) pre-equilibrated with 20 mM Tris-HCl (pH 7.4), 50 mM NaCl. NQO1 or NQO2 containing fractions were pooled, concentrated, frozen in liquid N_2_ and stored at −80 °C.

### Purification of CBR3

BL21(DE3) were transformed with pNIC28Bsa4-CBR3. Cells were grown in LB medium supplemented with 50 µg/ml kanamycin at 37 °C until they reached OD_600_ 0.6. Protein expression was induced by the addition of 1mM IPTG for 3 h. Cells were harvested by centrifugation at 5000 *g* for 10 minutes, and resuspended in 20 mM sodium dihydrogen phosphate pH 7.4, 20 mM Imidazole, 150 mM NaCl, 0.26 mM PMSF, 1 mM Benzamidine, 1.4 µg/ml Pepstatin. Cells were disrupted by the addition of 1 mg/ml lysozyme followed by rolling at 4°C for 30 mins, and sonication (40% amp, 30sec pulses for 7.5 minutes). The lysed cells were centrifuged at 18000 rpm for 45 mins at 4 °C to remove cellular debris. The supernatant was applied to a HisTrapHP column pre equilibrated in 20 mM sodium dihydrogen phosphate pH 7.4, 20 mM Imidazole, 150 mM NaCl. His-TEV-CBR3 was eluted with a linear gradient to 400mM imidazole over 40 mls. Fractions containing His-TEV-CBR3 were pooled and incubated at room temperature for 3 hours with TEV protease. The cleaved sample was dialyzed overnight against 20 mM sodium dihydrogen phosphate pH 7.4, 150 mM NaCl, then re-applied to a HisTrapHP column to remove uncleaved protein and TEV protease. The flowthrough was collected, concentrated using a 10 kDa Amicon Ultra column (Millipore) and loaded onto a gel filtration column (Superdex 200, 10/300 GL, GE Healthcare) pre-equilibrated with 20 mM sodium dihydrogen phosphate pH 7.4, 50 mM NaCl. CBR3 containing fractions were pooled, concentrated, frozen in liquid N_2_ and stored at −80°C.

### Purification of PGDH and RBBP9

BL21(DE3) were transformed with pET28-PGDH or pET28-RBBP9 with a C-terminal 6xHis tag. Proteins were purified as for CBR3 with the following changes. After elution from the first HisTrapHP column, fractions containing PGDH-His or RBBP9-His were concentrated and loaded onto a gel filtration column (Superdex 200, 10/300 GL, GE Healthcare) pre-equilibrated with 20mM sodium dihydrogen phosphate pH 7.4, 50 mM NaCl. Fractions containing PGDH- His or RBBP9-His were pooled, concentrated, frozen in liquid N_2_ and stored at −80°C.

### Purification of NRas, KRas, HRas and RhoA

BL21(DE3) transformed with pET28-MHL NRASA, pET28-MHL KRASB (H61), pET28-HRas or pNIC28-Bsa4 RhoA were induced and purified as for CBR3 with the following changes; Cells were resuspended in 20 mM Tris-HCl pH 7.4, 20 mM Imidazole, 150 mM NaCl, 0.26 mM PMSF, 1 mM Benzamidine, 1 µg/ml Pepstatin. TEV cleaved proteins were dialyzed against 20 mM Tris pH 7.4, 150 mM NaCl. The Superdex 200 10/300 GL gel filtration column was pre-equilibrated in 20 mM Tris pH 7.4, 50 mM NaCl.

### Purification of mammalian 20S proteasomes

Purification of the rat 20S proteasome was performed as described previously (Moscovitz et al., 2015). In brief, rat livers were homogenized in buffer containing 20 mM Tris-HCl pH 7.5, 1 mM EDTA, 1 mM DTT and 250 mM sucrose. The extract was subjected to centrifugation at 1,000 *g* for 15 min. The supernatant was then diluted to 400 ml to a final concentration of 0.5 M NaCl and 1 mM DTT and subjected to ultracentrifugation for 2.2 hours at 145,000 *g*. The supernatant was centrifuged again at 150,000 *g* for 6 hours. The pellet containing the proteasomes was resuspended in 20 mM Tris-HCl pH 7.5 and loaded onto 1.8 L Sepharose 4B resin. Fractions containing the 20S proteasome were identified by their ability to hydrolyze the fluorogenic peptide suc-LLVY-AMC, in the presence of 0.02% SDS. Proteasome-containing fractions were then combined and loaded onto four successive anion exchange columns: Source Q15, HiTrap DEAE FF and Mono Q 5/50 GL (GE Healthcare). Elution was performed with a 0–1-M NaCl gradient. Active fractions were combined, and buffer exchanged to 10 mM phosphate buffer pH 7.4 containing 10 mM MgCl2 using 10 kD Vivaspin 20 ml columns (GE Healthcare). Samples were then loaded onto a CHT ceramic hydroxyapatite column (Bio-Rad Laboratories Inc.); a linear gradient of 10–400 mM phosphate buffer was used for elution. The purified 20S proteasomes were analyzed by SDS–PAGE, activity assays and MS analysis.

### Purification of yeast 20S proteasomes

*S. cerevisiae* expressing FLAG-tagged 20S proteasome (Pre1) were grown in 4×700 ml YPD medium overnight at 30 °C. Cells were harvested at 5000 *g* for 20 minutes, the pellets rinsed in 10 ml water and centrifuged again at 5000 *g* for 20 minutes. The pellet was resuspended in 100 ml lysis buffer containing 50 mM Tris-HCl pH 7.5, 150 mM NaCl, 10% glycerol, 5 mM MgCl_2_, 1 mM PMSF, protease inhibitor cocktail (Complete, Roche), 250 U Benzonase (Millipore). Cells were lysed using a glass bead beater, pre-chilled with 50% glycerol and dry ice, with 1 minute pulses for 7 minutes total. The lysed cells were separated from the glass beads, and centrifuged at 35,000 *g* for 20 minutes at 4 °C to remove cell debris. The supernatant was collected, and incubated with 2 ml anti-FLAG M2 affinity gel (Sigma), pre rinsed with sequential washes of lysis buffer, Glycine pH 3.5 and lysis buffer, for 1.5 hours at 4 °C while gently rotating. The beads were collected, washed sequentially with lysis buffer containing 0.2 % NP40, lysis buffer, and lysis buffer containing 500 mM NaCl. The last wash was incubated on the beads for 1 hour at 4 °C, followed by a final wash in lysis buffer. 20S proteasomes were eluted using 500 mg/ml FLAG peptide in lysis buffer containing 15 % glycerol.

### Purification of Archaea 20S proteasomes

The α and β subunits of *T. acidophilum* 20S proteasome were expressed as separate fusion proteins with a TEV-cleavable His tag (α) or with a NusA-His tag (β) in BL21 (DE3) cells. Expression of both subunits was induced with the addition of 1mM IPTG, 37 °C for 3 hours (α) or for 5 hours (β) at 37 °C. Cells were collected by centrifugation at 5000 *g* for 20 min. Cells were lysed by sonication in 50 mM sodium phosphate buffer pH 8.0, supplemented with protease inhibitors (0.5 mM benzamidine, 0.1 mg/ml pepstatin A and 0.1 µM PMSF), 0.88 mg/ml lysozyme, and 250 U Benzonase (Millipore). After centrifugation at 40,000 *g* for 30 minutes, the supernatant was loaded onto a HisTrap FF (GE Healthcare) pre-equilibrated in 50 mM sodium phosphate buffer pH 8.0, 200 mM NaCl, 10 mM imidazole. The α and β subunits were eluted in 100 mM sodium phosphate buffer pH 7.8, 300 mM Imidazole. The fractions containing the fusion protein were pooled and dialyzed overnight with TEV protease against 50 mM Tris pH 7.4, 1 mM EDTA and 2 mM DTT. Following the overnight TEV cleavage, the α and β subunits were loaded onto a HisTrap FF column and flow through fractions were collected. The full proteasome (α_7_β_7_β_7_α_7_) was assembled by mixing a slight molar excess of α subunit over β subunit, and incubated at 37 °C for 6 hours. The mixture was then concentrated to 0.5 ml and incubated overnight at 37 °C. The assembled 20S proteasome complex was loaded onto a Superdex 200 10/300 GL (GE Healthcare) pre-equilibrated in 50 mM sodium phosphate buffer pH 7.5, 200 mM NaCl.

#### Proteasome degradation assays

To monitor the ability of proteins to regulate the activity of the 20S proteasome *in vitro*, 10 µM of the CCRs or MG132 were pre-incubated with 0.1 µM of the 20S proteasome for 30 minutes on ice in 50 mM HEPES pH 7.5. To initiate the assay, α-syn was added to 1 µM, and the reaction mixtures were incubated at 37 °C. For experiments using yeast or archaea 20S proteasomes, the experiments were performed at 25 °C. 10 µl samples were taken every 30 minutes for 120 minutes, quenched by the addition of reducing sample buffer and snap frozen in liquid N_2_. After all time points were collected, the samples were thawed, boiled for 5 minutes, and loaded onto a 15 % SDS-PAGE gel. Gels were stained with Commassie brilliant blue, and changes in the level of α-syn were quantified by band densitometry using ImageJ, normalized to T0, and plotted using Graphpad Prism.

Degradation assays using OxCaM were performed in the same way with minor modifications. OxCaM was added to a final concentration of 2.5 µM, and time points were collected every hour for 4 hours. Assays performed with RBBP9, NRas, KRas, HRas and RhoA were analyzed by western blot with anti-calmodulin antibody, due to these CCRs being the same size as OxCaM.

#### Proteasome activity assays

Proteasome activity assays were performed as previously described (Moscovitz et al., 2015). In brief, between 0.1-0.3 µM 20S proteasomes were incubated either alone or with 10 µM MG132 or CCRs in 25 mM HEPES pH 7.5 for 30 mins on ice. 4 µl of the mixture was combined with 40 µl 25 mM HEPES pH 7.5 containing 100 uM Suc-LLVY-AMC and 0.02 % SDS. Samples were incubated at 30 °C for 30 min in the dark. To stop the reaction, 200 µl 1 % SDS was added to the mixture. The fluorescence of hydrolyzed AMC groups was measured with a microplate reader (Infinite 200, Tecan Group), using an excitation filter of 380 nm and an emission filter of 460 nm.

#### Native mass spectrometry analysis

Nanoflow electrospray ionization MS and tandem MS experiments were conducted under non-denaturing conditions on a Q-Exactive Plus Orbitrap EMR (ThermoFisher Scientific). Before MS analysis, 20 µl of up to 100 mM sample was buffer exchanged into 0.5–1 M ammonium acetate pH 7.5, using Bio-Spin columns (Bio-Rad). Assays were performed in positive ion mode and conditions were optimized to enable the ionization and removal of adducts, without disrupting the non-covalent interactions of the proteins tested. In MS/MS experiments, the relevant m/z values were isolated and argon gas was admitted to the collision cell. Spectra are shown without smoothing or background subtraction. Typically, aliquots of 2 µl of sample were electrosprayed from gold-coated borosilicate capillaries prepared in-house. The following experimental conditions were used on the Q-Exactive Plus Orbitrap EMR: capillary voltage 1.7 kV, MS spectra were recorded at low resolution (5000), and the HCD cell voltage was set to 20– 50 V, at trapping gas pressure setting of 3.9. For MS/MS analyses, a wide isolation window of ±2000 m/z around the most intense charge state of the 20S proteasome (around 12,000 m/z) was set in the quadrupole, allowing the transmission of only high m/z species. Transmitted ions were subjected to collision induced dissociation in the HCD cell, at an accelerating voltage of 200 V, and the trapping gas pressure was set to 1.5.

#### Immunoprecipitation

HEK293T cells stably expressing the FLAG-β_4_ proteasome subunit were plated in four 15-cm dishes, at a density of 1.5×10^6^ cells per dish. Each plate was transfected with 20 µg of pCDF1- NQO2-HA, pCDF1-PGDH-HA, pCDF1-NRas-HA or pCDF1-RhoA-HA and grown for 48 hours. Cells were collected by trypsinization, combined, washed in PBS and resuspended in 1 ml lysis buffer (20 mM HEPES pH 7.4, 10 % glycerol, 10 mM NaCl, 3 mM MgCl_2_, 1 mM ATP) and protease inhibitors (1 mM PMSF, 1 mM Benzamidine, 1.4 µg/ml Pepstatin). Cells were incubated on ice for 15 min and homogenized in a glass-Teflon homogenizer for 40 strokes. Lysate was cleared by centrifugation at 18,000 *g* for 10 min at 4 °C. For IP using anti-FLAG affinity gel, 1 mg protein was diluted in 500 µl lysis buffer. NaCl concentration was adjusted to 150 mM, and rotated overnight at 4 °C in the presence of 45 µl anti-FLAG M2 affinity gel (Sigma). The following morning, beads were washed three times with lysis buffer containing 150 mM NaCl and boiled in 55 µl reducing sample buffer. For IP using anti-HA or anti-Rpn2 antibodies, 1 mg protein was diluted in 500 μl lysis buffer. NaCl concentration was adjusted to 150 mM. Proteins were pre-cleared using 40 μl of Protein G Sepharose (GE Healthcare), for 1 hour at 4 °C, at a gentle rotation. The beads were discarded and the lysate was rotated overnight at 4 °C in the presence of 9 μl anti-HA rabbit (ab9110, Abcam) or 9 μl anti-Rpn2 (PSMD1, ab140682, Abcam) antibody. The following morning, 45 μl Protein G Sepharose beads (GE Healthcare) were added, and lysate was rotated for 2 h at 4 °C. The beads were then washed three times in lysis buffer containing 150 mM NaCl and boiled in 55 μl protein sample buffer.

#### CCR Overexpression

A31N-ts20 BALB/c cells were transfected by electroporation using a NEPA21 electroporator (Nepa Gene Co., Ichikawa-City, Japan). For electroporation, 2#x00D7;10^6^ cells were mixed with 20 µg pCDF1 plasmid containing GFP, NQO2, CBR3 or PGDH, and transfected according to the manufacturers instructions, using a poring pulse of 125V for 5ms. After electroporation, cells were cultured at 32 °C for 24 hours. The growth medium was then replaced and the cells returned to 32 °C for 24 hours. Cells were transferred to 39 °C for 24 hours, or left at 32 °C as indicated. Seventy-two hours post transfection, the cells were collected and lysed in modified RIPA buffer containing 50 mM HEPES pH 7.5, 150 mM NaCl, 1% NP-40, 0.25% Na-deoxycholate, 0.26 mM PMSF, 1 mM Benzamidine, 1.4 µg/ml Pepstatin, 4 mM Na-pyrophosphate, 4 mM β-glycerophosphate, 5 mM Na-orthovanadate. Cellular debris was removed by centrifugation, and the supernatant was collected. Total protein concentration was estimated by Bradford assay. For western blot analysis, 30 µg total protein was loaded for each sample.

#### Cell fractionation

MCF10A cells from a 6 cm tissue culture dish were resuspended in 70 µl hypotonic buffer, containing 10 mM HEPES pH 7.4, 10 mM KCl, 1.5 mM MgCl2, 0.5 % NP-40, 4 mM Na-pyrophosphate, 4 mM β-glycerophosphate, 5 mM Na-orthovanadate, 0.26 mM PMSF, 1 mM benzamidine, 1.4 µg/ml pepstatin, and incubated on ice for 10 minutes. 5 µl of 10% NP-40 was added and cells were vortexed for 10 s. The cytosolic fraction was separated from the nuclei by centrifugation at 2500 rpm for 4 minutes and further clarified by centrifugation at 10,000 *g* for 10 minutes. The nuclear pellet was washed in 0.1 ml hypotonic buffer and nuclei were pelleted at 2500 rpm for 4 minutes. Nuclei were then resuspended in 35 µl hypertonic buffer, containing 20 mM HEPES pH 7.6, 100 mM NaCl, 300 mM sucrose, 3 mM MgCl_2_, 1 mM CaCl_2_, 0.5 % TritonX-100, 0.26 mM PMSF, 1 mM benzamidine, 1.4 µg/ml pepstatin, and incubated on ice for 3 minutes. The nucleoplasmic fraction was separated by centrifugation at 5,000 *g* for 5 minutes. The pellet was resuspended in 28 µl hypertonic buffer and incubated at room temperature for 15 min with 1000 gel units of micrococcal nuclease (NEB). Chromatin-bound proteins were released from the DNA by addition of 7 µl of 1 M ammonium sulfate on ice for 5 minutes, followed by centrifugation at 5,000 *g* for 5 minutes. Nucleoplasmic and chromatin-bound fractions were combined. 30 µg of cytoplasmic and 20 µg total nuclear fractions were loaded onto 12 % SDS-PAGE gels, followed by western blot analysis.

#### Western blot

After separation of samples on SDS-PAGE, proteins were transferred to 0.45 µm Immobilon-P PVDF membranes (Millipore) pre activated in methanol, in standard Tris-Glycine transfer buffer (pH 8.3) supplemented with 20% methanol for 2.5 hours at 400 mA. Membranes were blocked in 5 % skim milk powder in TBS-T for 1 hour, followed by incubation with appropriate primary antibodies on an orbital shaker at 4 °C overnight. Membranes were rinsed thoroughly in TBS-T, followed by incubation with appropriate secondary HRP conjugated antibodies for 1 hour on an orbital shaker at room temperature. Membranes were rinsed thoroughly and developed using WesternBright ECL (Advansta) in a MyECL Imager (Thermo Scientific) according to the manufacturer’s instructions.

Primary antibodies used for western blots include anti-calmodulin (1:1000, ab105498, Abcam) anti-HA rabbit (1:6000, ab9110, Abcam), anti-HA mouse (1:1000, ab18181, Abcam), anti-PSMD1 (1:1000, ab2941, Abcam), anti-PSMA1 (1:1000, ab140499, Abcam), anti-FLAG (1:2500, F3165, Sigma), anti-GFP (1:2500, ab290, Abcam), anti-NQO2 (1:500, sc271665, Santa Cruz), anti-CBR3 (1:1000, 15619-1-AP, Proteintech), anti-PGDH (1:200, sc271418, Santa Cruz), anti-p53 HRP (1:2500, HAF1355, Biotest), anti-αsyn (1:500, ab51252, Abcam), anti-GAPDH (1:1000, MAB374, Millipore), anit-Ubiquitin (1:1000, PW0930, Enzo), anti-p21 (1:1000, ab109199, Abcam), anti-Tubulin (1:10,000, ab184613, Abcam). Anti-Nrf2 (1:500, ab137500, Abcam), anti-HistoneH3 (1:1000, ab24834, Abcam), anti-NQO1 (1:1000, ab28947, Abcam), anti-NRas (1:200, sc31, Santa Cruz), anti-DJ-1 (1:10,000, ab76008, Abcam), anti-RhoA (1:200, sc418, Santa Cruz), anti-RBBP9 (1:1000, 12230-2-AP, Proteintech).

Secondary antibodies used for western blots include Goat anti-mouse IgG-HRP (1:10,000, 115- 035-003, Jackson) and Goat anti-rabbit IgG-HRP (1:10,000, 111-035-003, Jackson).

#### Quantification and Statistical Analysis

Where indicated in the figure legends, at least three independent biological replicates were performed. The n number of experiments and the details of the statistical analysis are described in the figure legends. Students t-tests were performed to measure statistical significance, which is defined as * p < 0.05, ** p < 0.01, *** p < 0.001, **** p < 0.0001. All error bars correspond to S.E.M. as indicated in the figure legends. Statistical analysis was performed using GraphPad Prism.

## Supplemental Information

**Supplementary Figure 1:**
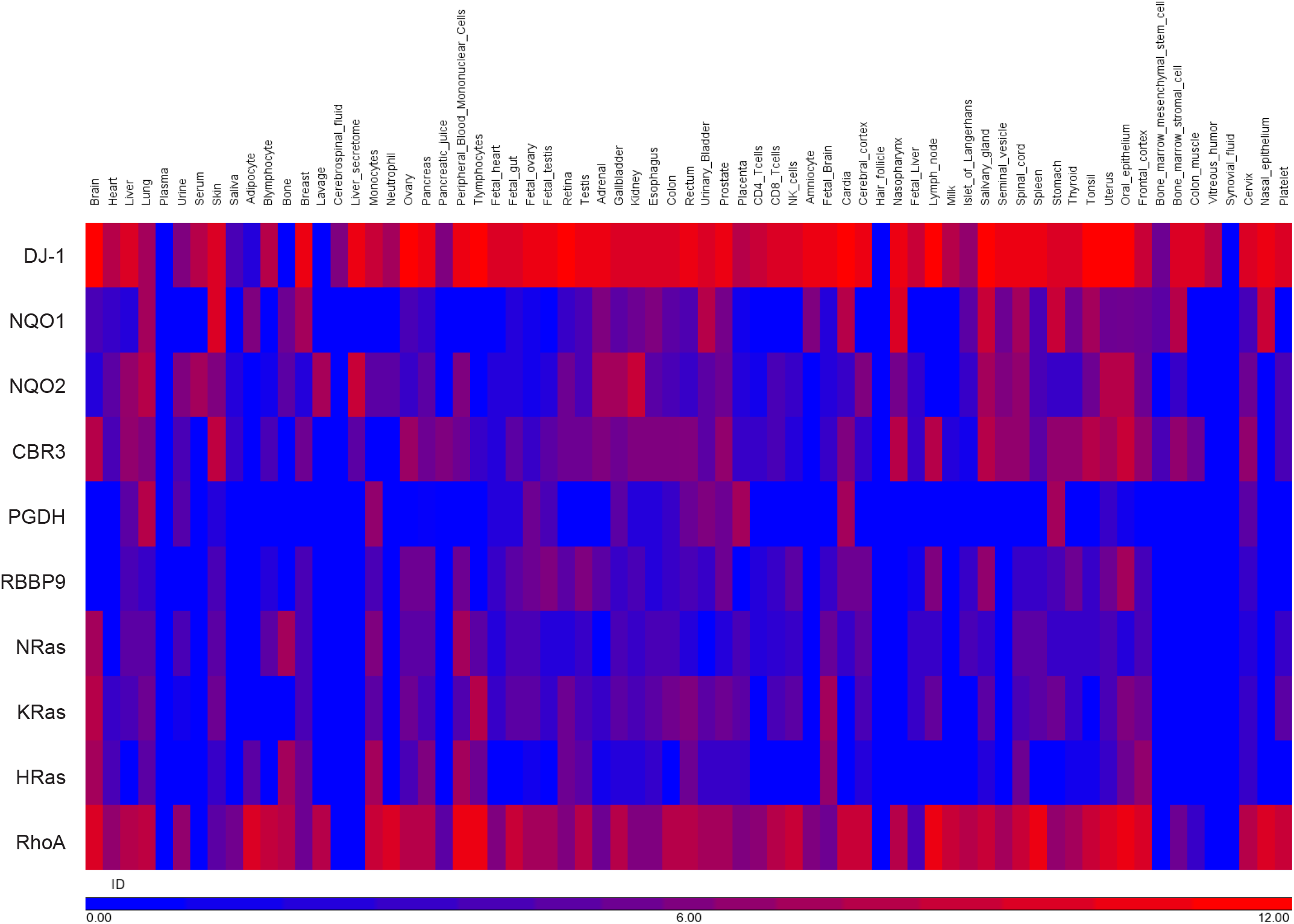
Tissue wide protein expression of the CCRs. Estimated protein expression levels of the CCRs analyzed in this study in various human tissues, taken from the Human Integrated Protein Expression Database (HIPED) (Fishilevich et al., 2016). DJ-1 and RhoA display the broadest expression across all tissues. Expression levels are presented in log2(ppm). Figure was prepared using Partek Genomics Suite 7.18.

**Supplementary Figure 2:**
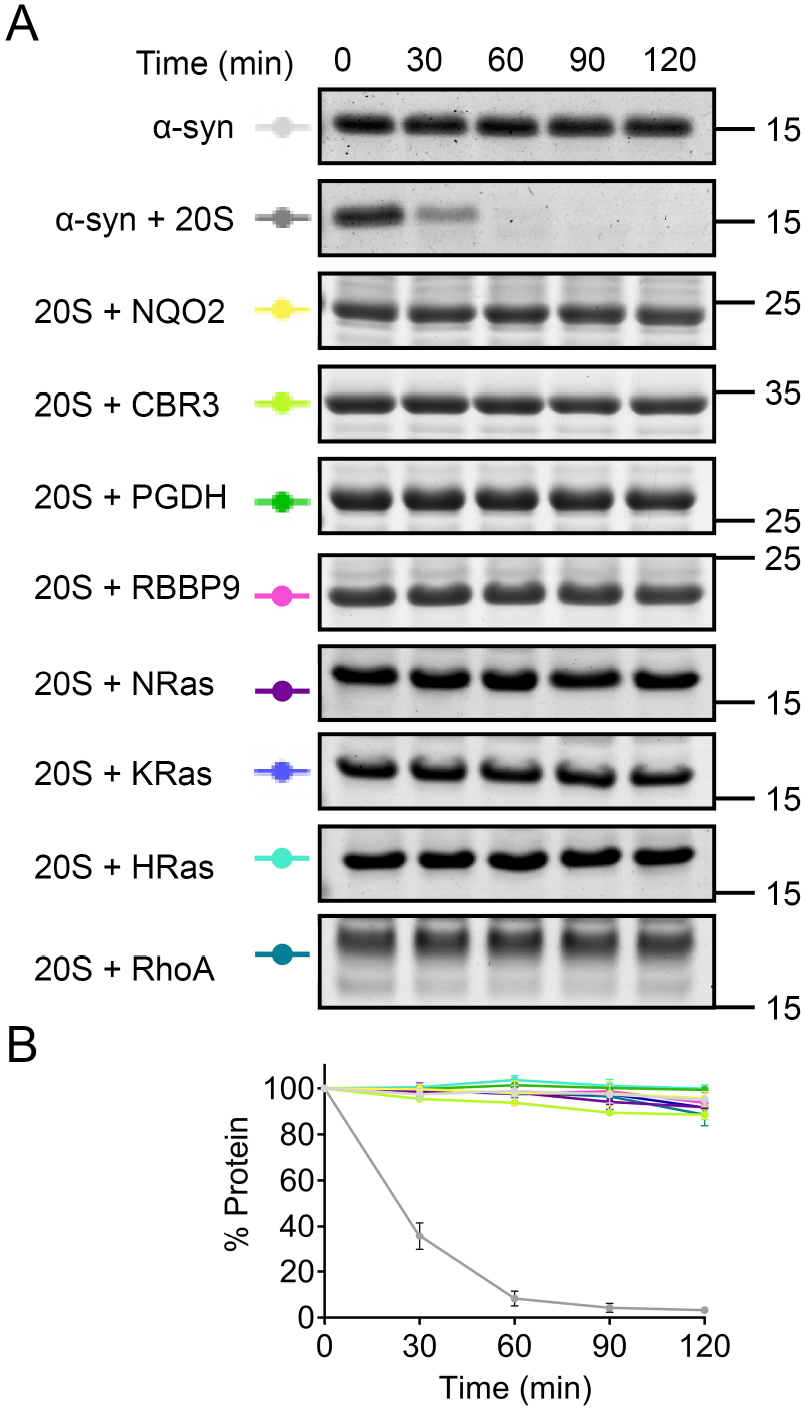
CCRs are not degraded by the 20S proteasome. (A) *In vitro* degradation assays of each CCR with 20S proteasome in the absence of α-syn. As controls, α-syn alone and in the presence of 20S proteasome (top two panels) is included to ensure active 20S proteasome. (B) Quantification of α-syn (from control panels in A) or each CCR from three independent experiments. Error bars represent S.E.M.

**Supplementary Figure 3:**
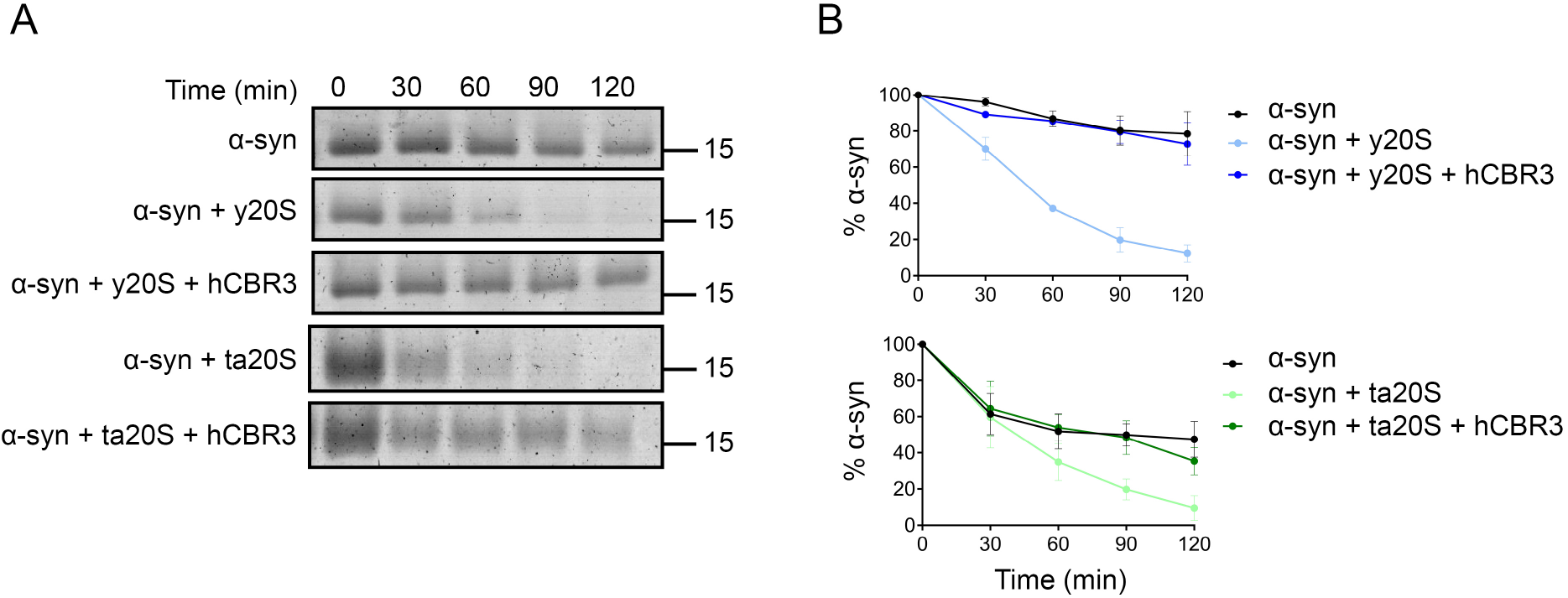
The inhibitory activity of CBR3 on 20S proteasome substrate degradation is conserved across evolution. (A) In vitro degradation assays using model substrate α-syn were performed using human CBR3 (hCBR3) with 20S proteasomes from yeast (*S. cerevisiae*, y20S) or archaea (*T. acidophilum*, ta20S). (B) Quantification of three independent experiments is shown, error bars represent S.E.M. (C) Weblogo derived from MSA analysis of CCR homologues, conserved N-terminal motif is seen between residues 16-24.

**Supplementary Figure 4:**
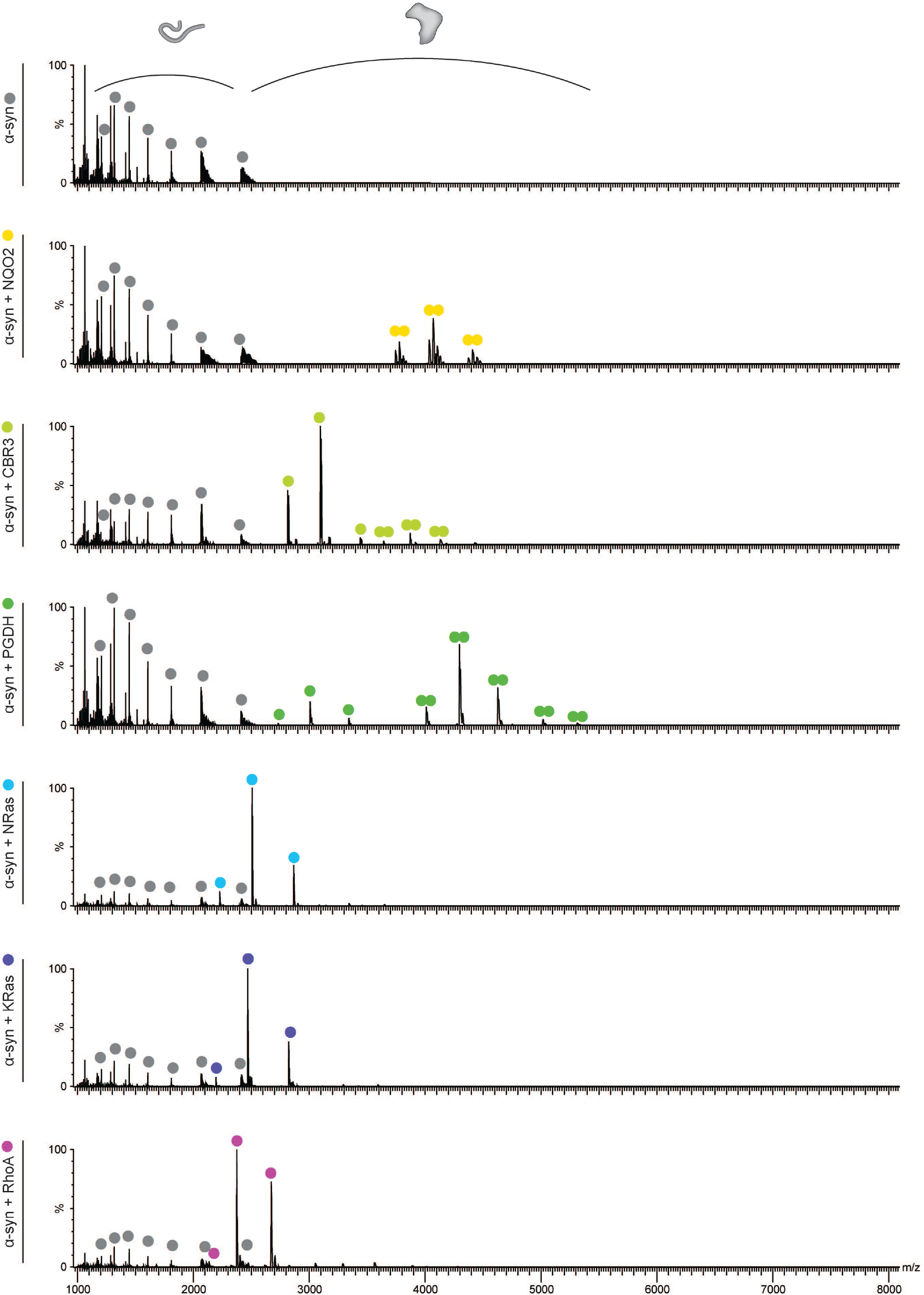
Native MS does not detect any interaction between substrates and CCRs. α-syn was analyzed by native mass spectrometry either alone (top panel) or in the presence of each of the CCRs. The charge series corresponding to α-syn was measured in each spectrum (gray balls). Each of the CCRs were detected in their respective spectra (NQO2 - yellow balls, CBR3 – lime balls, PGDH – green balls, NRas – dark purple balls, KRas – dark blue balls, RhoA – teal balls). No larger molecular weight complexes were detected in any of the spectra, indicating that α-syn does not bind to any of the CCRs.

**Supplementary Table 1:** Predicted and measured masses of 20S proteasome subunits, CCRs and α-syn detected by MS/MS.

| Protein name | Uniprot Accession Number | Modifications | Theoretical Monomeric Mass (Da) | Measured Mass (Da) |
| --- | --- | --- | --- | --- |
| PSMA2 | P25787 | Δmet, acetyl | 25,838 | 25,838 +/- 1.4 |
| PSMA4 | P25789 | Δmet, acetyl | 29,409 | 29,409 +/- 1.1 |
| PSMA5 | P28066 | acetyl | 26,453 | 26,453 +/- 0.7 |
| PSMA6 | P60900 | Δmet, acetyl | 27,311 | 27,311 +/- 1.2 |
| PSMA7 | O14818 | Δmet, acetyl | 27,767/27,804 | 27,609 +/- 1.1 |
| NQO2 | P16083 | + Zn<sup>2+</sup> | 26,176 | 26,174 +/- 1.1 |
| CBR3 | O75828 | +Nterm S | 30,937 | 30,937 +/- 1.1 |
| PGDH | P15428 | 6His | 30,042 | 30,041 +/- 0.6 |
| RBBP9 | O75884 | Δmet, 6His | 21,934 | 21,935 +/- 0.7 |
| NRas | P01111 | 1-172, +Nterm G | 19,604 | 19,606 +/- 0.4 |
| KRas | P01116 | 1-169, +Nterm G | 19,303 | 19,304 +/- 0.3 |
| HRas | P01112 | 1-172, +Nterm G | 19,764 | 19,764 +/- 0.3 |
| RhoA | P61586 | 1-184, +Nterm S | 20,898 | 20,897 +/- 0.6 |
| α-syn | P37840 | - | 14,460 | 14460 +/- 1.0 |

Modifications: $\Delta$ met (loss of N-terminal Met), acetyl (acetylation), Zn<sup>2+</sup> (Zn ion bound as cofactor), Nterm SG (residual N-terminal Ser/Gly after cleavage of affinity tag), 6His (6x His tag was not cleaved during purification), 1-1xx (truncation of full length protein to aid solubility).

